# Community coalescence reveals strong selection and coexistence within species in complex microbial communities

**DOI:** 10.1101/2025.11.06.687011

**Authors:** Sophie Jean Walton, Qing Xu, Richa Sharma, Hannah R. Gellert, Chih-Fu Yeh, Jonas Cremer, Katherine S. Xue, Dmitri A. Petrov, Benjamin H. Good

## Abstract

Complex microbial ecosystems harbor extensive intra-species diversity, but the fitness consequences of this genetic variation are poorly understood in community settings. Here we address this question by competing in vitro gut communities derived from different human donors, revealing the emergent fitness differences between conspecific strains as they competed within larger communities. Most pairs of strains experienced strong and context-dependent selection, even when their parent communities were originally selected in the same nutrient environment. However, these fitness differences typically attenuated over time due to biotic interactions within the community, leading to extended coexistence within many species, and competitive exclusion in others. These results support the view that conspecific strains can fulfill distinct ecological roles when competing within a diverse community, even when their genomic diversity exhibits the hallmarks of a single biological species.

## Introduction

Large microbial communities, from the soil to the human gut, harbor diversity over multiple spatial and taxonomic scales (*1–8)*. Their vast species diversity is accompanied by extensive genetic variation within each of their component species, with strains from different local communities typically varying at thousands of genomic loci (*6, 9–19)*. On longer timescales, these conspecific strains compete with each other as part of a broader global population (*9, 20–24)*. They also compete on shorter timescales when new strains of an existing species invade from outside their local community. Examples of these inter-strain competitions are readily observed in longitudinal sequencing of microbial communities (*7, 25–30)*, and are conjectured to play an important role in driving their short-term (*9, 21, 26, 31)* and long-term (*9, 14, 21, 23)* evolution. However, the evolutionary forces that govern such local strain competitions remain poorly understood empirically. A central challenge lies in understanding how natural selection acts on strains of the same species that compete within larger, species-rich communities.

One prominent view is that selection on genetically diverged strains should be very strong, with one strain rapidly dominating over the other. This scenario is motivated by previous work showing that circulating strains often vary in many fitness-relevant traits, including resource utilization (*15, 19)*, inter-bacterial antagonism (*16)*, and phage susceptibility (*32, 33)*. The large number of such differences suggests that their cumulative effect on relative fitness could be large in any given local environment. On the other hand, ecological theory predicts that the assembly of highly diverse communities can select for strains that fulfill very similar functional roles within their local community context. This can lead to weak selection or effective neutrality among co-occurring strains that are otherwise phenotypically distinct (*34–40)*. Signatures of this emergent reduced selection have previously been observed at the species level (*41)*, suggesting that it might naturally extend to strain-level competition at finer genetic scales.

A third possibility is that strains can coexist with each other despite strong selective differences by competing for distinct ecological niches (*42)*, with their extensive phenotypic differences allowing them to function more like different species in their local community context. This niche partitioning model is motivated by empirical observations showing that multiple strains of the same species can sometimes persist within a local community for extended periods of time (*26– 30, 43–46)*, even when their historical patterns of gene flow are more consistent with a single biological species (*20, 21, 23, 47)*. However, the strength of niche partitioning within species is still debated. Theoretical models predict that competition with other species can severely limit the realized niche of a given species (*40, 48–52)*, which could make it more difficult for conspecific strains to engage in niche partitioning when competing within a larger community. Understanding how coexistence emerges in these species-rich settings is a critical open problem, which has important implications for the design of microbial therapeutics (*53–58)* and the dynamics of strain transmission across hosts (*25, 31, 46, 59–61)*.

Distinguishing these scenarios can be challenging in natural microbial communities due to the confounding effects of spatial structure (*62, 63)*, migration (*25, 59, 60)*, and stochastic environmental variation (*7, 25, 26, 62, 64)*. In vitro systems provide a promising alternative, by enabling quantitative measurements of microbial competition in controlled laboratory conditions. In particular, several recent studies have shown that anaerobic passaging of human fecal samples can yield highly diverse, stable, and reproducible microbial communities that can be serially propagated in liquid culture for extend periods of time (*41, 65–68)*. We reasoned that these stool-derived communities could provide a natural platform for investigating strain-level competition within a complex community setting, by leveraging the extensive strain-level variation that already exists within and among different people.

Here we illustrate this approach by pairing strain-resolved metagenomics with serially passaged stool communities derived from different human donors. By tracking whole-community mixtures at high temporal and genetic resolution, we measured the emergent fitness differences between multiple pairs of conspecific strains as they competed within larger, species-rich communities. These community-scale fitness assays provide a window into the eco-evolutionary forces acting on circulating strains within complex microbial communities.

## Results

### In vitro communities assembled from human stool samples contain a range of strain-level diversity

To study strain-level competition in a complex community setting, we derived in vitro communities from frozen stool samples from three healthy human donors (Methods). For each subject, we used aliquots from a single homogenized sample to inoculate three replicate communities in each of two distinct media (mBHI and mGAM) that promote the growth of different species (*65–68)*. We propagated each community via 1:200 dilutions into fresh media every 48 hours for five total passages and froze the resulting communities in glycerol for future use. We then performed shotgun metagenomic sequencing on each timepoint as well as the original stool inoculum, and used a read-mapping approach to characterize species relative abundances and the genetic variation within each species (Methods). Consistent with previous results (*41, 65–68)*, we found that the in vitro communities rapidly converged to diverse steady states within ∼2-3 passages (Figs. 1A,B, S1,S2; Methods) with ∼50 species present above a relative abundance of 0.1% (IQR=46-53). While the final species composition differed from the original stool inocula (Figs. 1C and S2; p<0.001, permutation test in the difference in median Jensen-Shannon distance, JSD), the overall species richness remained high and contained prominent gut bacterial families, such as *Bacteroidaceae* (Figs. 1A,B and S1). The species composition of the assembled communities was also specific to the donor and media in which they were grown (Fig. 1C, p<0.001), though many individual species were also shared (Fig. S2; 65–68).

**Fig. 1:**
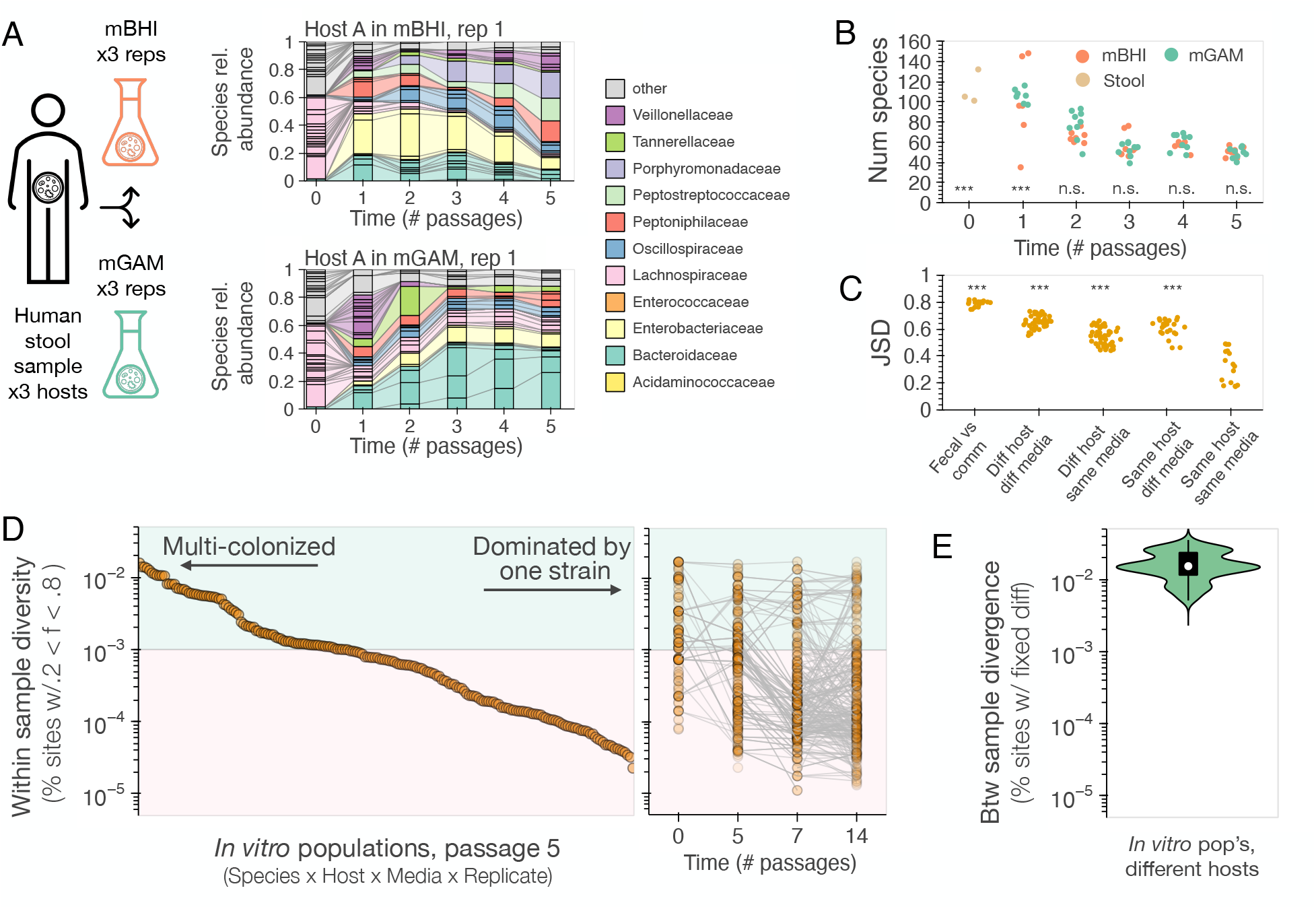
Assembly of in vitro communities from human stool samples yields species-rich communities with a range of strain-level variation. **(A)** Stool samples from three healthy adults were serially passaged in mBHI or mGAM media every 48 hrs in 1:200 dilutions (Methods). Two example communities are shown. Bars denote species relative abundances estimated from metagenomic sequencing (Methods), colored by bacterial family. **(B)** Species richness over time, estimated as the total number of species with relative abundance >0.1%. Asterisks show permutation tests on the difference in medians compared to passage 5 (***=p<0.001; n.s.=p>0.05). **(C)** Jensen-Shannon distance (JSD) between communities at the end of passage 5, demonstrating that community composition is host- and media-specific (permutation test on the difference in medians compared to biological replicates, ***=p<0.001). **(D)** Intra-species genetic diversity, estimated from the fraction of polymorphic sites per sample, for all species with sufficient sequencing coverage (Methods). Left panel shows the distribution of diversity at passage 5, while right panel shows the evolution of this metric over an additional 9 passages, including a freeze-thaw cycle after passage 5. Adjacent timepoints from the same population are connected by solid lines to aid in visualization. Many in vitro populations exhibit high levels of genetic diversity, comparable to the typical genetic differences between strains in different hosts **(E)**. Horizontal shading in D denotes the threshold used to classify populations as being dominated by a single vs multiple colonizing strains (Methods).

To investigate the genetic diversity within species, we used a reference-based approach to identify single nucleotide variants (SNVs) within all species with >5x coverage at a given timepoint (Methods). Similar to observations in human stool samples (*9, 69)*, we found that our in vitro populations varied widely in the total number SNVs that were present at intermediate frequencies (Fig. 1D). Some species’ populations had just a handful of polymorphisms, while others had tens of thousands – comparable to the typical genetic differences between strains in different hosts (Figs. 1E and S3). Previous work has shown that these high-diversity populations correspond to cases where the host was originally colonized by two or more genetically diverged strains of the same species (*9, 69)*. By contrast, the lower diversity populations correspond to cases where the species is dominated by a single resident strain, whose genotype can be reliably inferred from the consensus allele in the population.

Using previously established thresholds based on the inter-subject divergence in Fig. 1E (*9)*, we classified each population as single- or multi-colonized if its within-sample diversity was less than or greater than 0.1% (Methods). We found that around half of the assessed populations at passage 5 showed evidence of multi-colonization by this criterion (n=82/200, 41%; Fig. 1D). We confirmed that these signatures were not driven by cross-contamination by examining patterns of SNV sharing across samples from different hosts (Fig. S3; Methods; 9, 10, 25, 70). Intra-species diversity was also consistent across independent replicates from the same inoculum: of the 34 populations with at least one multi-colonized replicate, a total of 21 (62%) had another replicate that was also classified as multi-colonized, which was significantly higher than expected by chance (p<0.001; permutation test, Methods). Since any non-growing strains would be rapidly diluted out during community propagation, these observations indicate that multiple conspecific strains were able to grow and compete within our in vitro communities over multiple rounds of passaging.

To examine the persistence of these trends over longer timescales, we revived a subset of the frozen communities at timepoint 5 and passaged them for an additional 9 growth cycles (Fig. 1D). While intra-species diversity declined in many of the multi-colonized populations by passage 14 (n=22/30, 73%), a substantial number continued to exhibit high diversity at the end of this second round of passaging (n=8/30, 27%). These observations show that multiple conspecific strains can be maintained in serially passaged communities for extended periods of time, echoing previous results in other in vitro passaging systems (*45, 71)*.

### Community coalescence reveals strong selection on conspecific strains from different host communities

The signatures of multicolonization in Fig. 1D could be driven by nearly neutral competition within species, as well as stable coexistence through niche partitioning. To distinguish between these different scenarios, we sought to perform more controlled investigations of selection on con-specific strains without a prior history of co-occurrence. We therefore turned to whole-community competitions [also known as “community collisions” (*72)* or “community coalescence” (*73, 74)*], in which established communities are mixed together and propagated further over time. Community coalescence experiments have previously been used to study species-level behavior (*41, 75– 78)*, and are reminiscent of fecal microbiome transplants in therapeutic contexts (*27, 43)*. Here we leveraged the same idea to study the competition between strains of the same species derived from different host communities.

We implemented this approach by mixing pairs of assembled communities from passage 7 of our initial experiment in equal volumes, yielding a total of 6 initial inocula (3 pairwise mixtures in each of two assembly environments) and 6 autologous controls (Fig. 2A and S4-S7). We then split each inoculum into eight replicate collision experiments, four of which were passaged in mBHI and four passaged in mGAM, for 7 additional passages. We performed metagenomic sequencing on each community at all timepoints to monitor their species and strain dynamics over time. Since the assembled communities from different hosts shared 6-9 species at a relative abundance >1% (Fig. S2), we reasoned that this experimental design would allow us to measure competition between multiple strains in parallel, each of which was previously selected to be able to grow in at least one in vitro community.

**Fig. 2:**
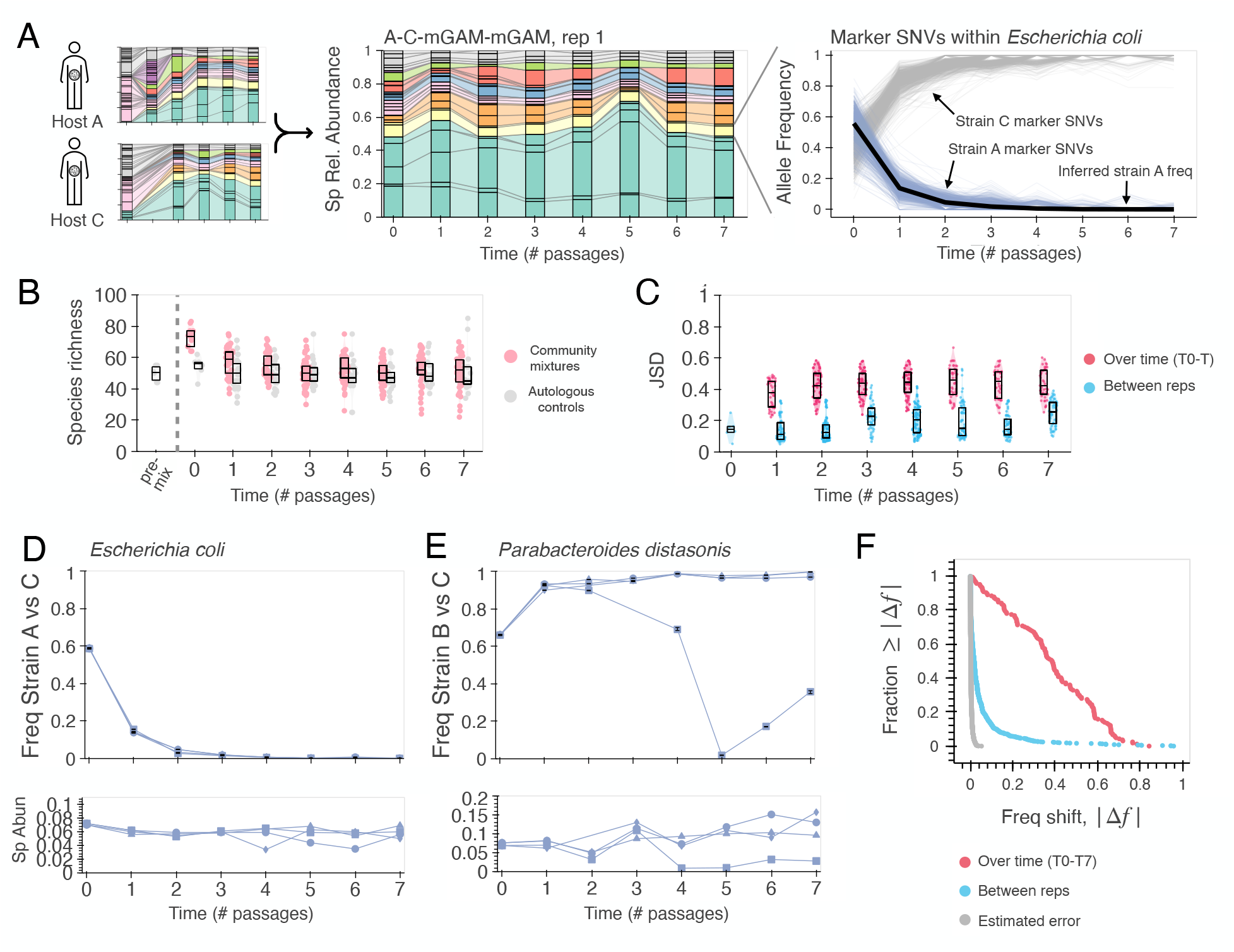
Community coalescence reveals strong selection on conspecific strains from different host communities. **(A)** In vitro communities from different subjects were mixed in equal ratios and passaged for 7 additional growth cycles. Left panels show species abundances over time for one example community, using the same color scheme as in Fig. 1A. Right panel shows the frequencies of marker SNVs within the *E. coli* sub-population of the same community. Thin lines represent the frequencies of the non-reference alleles at sites where the *E. coli* strains from the two parent communities differed from each other, while the heavier black line shows the inferred relative frequency of the strain from subject A (Methods). **(B)** Species richness of coalescing communities was elevated compared to autologous controls in the first two passages, but converged to similar levels at later timepoints. Boxes show the median and inter-quartile range at each timepoint. **(C)** Coalescing communities exhibit reproducible shifts in community composition over time. Points show the Jensen-Shannon distance (JSD) relative to the initial inocolum (pink) and between biological replicates (blue). **(D)** Inferred strain frequencies for all four replicates of the community collision in panel A (top), and the corresponding relative abundance of the entire species (bottom). Individual replicates are illustrated with distinct marker shapes. Error bars indicate approximate 95% confidence intervals in the inferred strain frequencies (Methods). (**E**) Analogous species and strain trajectories from a different example species where one replicate diverged from the others. In this example, the diverging strain frequencies (squares) were accompanied by a divergence in the relative abundance of the entire species. **(F)** Distribution of the net change in strain frequency over time across all 164 pairwise competitions (pink), compared to the differences between replicates (blue) or the inferred measurement error (grey, Methods). The temporal shifts in strain frequencies generally exceeded other sources of experimental noise.

Consistent with previous work (*41)*, we found that the species composition of each collision rapidly approached a new steady state within ∼ 3-4 passages (Figs. 2A-C, S8, and S9; Methods). These species compositions were highly reproducible across replicates and were specific to the initial inocula and the media in which they were competed (Figs. 2C, S10, p<0.001 permutation tests, Methods). As seen in previous work (*41)*, we found that many species in the initial community mixtures appeared to go extinct over the first few passages, in a manner suggestive of competitive exclusion (Figs. 2B, S11, S12). Across all collisions, the average species richness significantly declined relative to the richness of the initial mixture (p<0.001; permutation test in the difference in medians), and approached the steady-state richness of autologous control communities that were passaged for the same amount of time (p>0.05). These data suggest that species-level selection is strong during community coalescence, with some of the originally assembled species competing for the same ecological niches (*41, 75)*.

To measure the strain-level competition within species, we leveraged the fact that the dominant strains in each host typically varied at thousands of distinct marker SNVs (Figs. 1E, S13). By tracking the allele frequencies of these SNVs using the reference-based pipeline above, one can infer the relative frequencies of the two strains over time (Fig. 2A, right) with an estimated resolution that is typically less than ∼1% (IQR=0.1-0.49%; Figs. 2F, S13, and S14; Methods). For simplicity, we restricted our attention to the mono-dominated samples in Fig. 1D where the marker SNVs for each strain could be inferred with a high degree of confidence (Fig. S13; Methods). Applying these filters yielded a total of 61 pairwise strain competitions across 13 bacterial species (Table S5).

We found that these conspecific strains often underwent large shifts in relative frequency over the first few passages of the collision (Fig. 2D-F). The magnitudes of these shifts were significantly larger than expected from measurement error or population bottlenecks during passaging (p<0.001, permutation test; Fig. 2F, Methods), suggesting that they were instead driven by differential growth within the in vitro community environment. Strain dynamics were also highly similar across replicate competitions, with most (88%) replicates differing by less than 10% from each other (Fig. 2F). As a result, the overall direction of the frequency shift between timepoints 0 and 7 was consistent across replicates in ∼70% of cases (Fig. S15), and was significantly higher than expected by chance (p<0.001, Methods). The magnitudes and reproducibility of these shifts imply that the intra-species dynamics were driven by strong and deterministic selection – even when the relative abundance of their focal species exhibited smaller shifts over time (Fig. S16).

While most competitions were consistent across replicates, we also observed some cases where the strain-level trajectories were more variable. For example, in the *Parabacteroides distasonis* competition in Fig. 2E, one replicate trajectory underwent a large reversal in frequency, while the other three replicates remained highly similar to each other. As above, the magnitude of this difference was far larger than expected from measurement error or demographic noise (p<0.001, Methods), suggesting that it was likely driven by idiosyncratic differences in the selective landscape within that particular replicate community. Consistent with this hypothesis, we found that the species abundance of *P. distasonis* also systematically diverged in this replicate (Fig. 2E, bottom), suggesting that it experienced a different growth environment possibly due to subtle shifts in the local biotic landscape. This same pattern also held more broadly, with a statistically significant correlation between species- and strain-level deviations between replicates (Fig. S17, p<0.001, Pearson’s correlation). These observations further highlight that conspecific strains can experience strong selective differences when competing within a larger community, even in communities that were assembled in the same abiotic environment.

### Selection shifts over time, allowing for strain coexistence

To more precisely quantify the selective forces acting on the conspecific strains in Fig. 2, we estimated their standard relative growth rate,

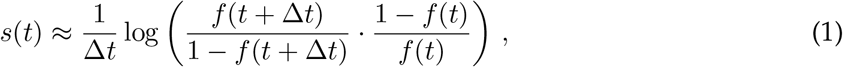

from their frequency trajectory *f* (*t*); this provides an operational definition of their relative fitness in the interval between *t* and *t* + 6.*t*. This difference in per capita growth rates can be computed for any two lineages within a community, regardless of whether they are competing within the same ecological niche. In the latter case, *s*(*t*) reduces to the standard competitive fitness from population genetics. In more general contexts, Eq. (1) will also account for ecological feedbacks within the community (frequency-dependent selection; 79), including any ecological interactions between the focal strains themselves.

Using this metric, we found that most pairs of strains exhibited large fitness differences during the early stages of competition (Figs. 3B and S18). The typical selection coefficients between passages 1 and 2 ranged from 0.95-2.66 per passage (IQR), comparable to the largest strain-level fitness differences that have been inferred from in vivo microbiome sequencing (*9, 12, 26, 80–83)*. If selection were constant over time, these relative fitness differences are large enough that the fitter strain would be expected to dominate the population over the next several passages.

**Fig. 3:**
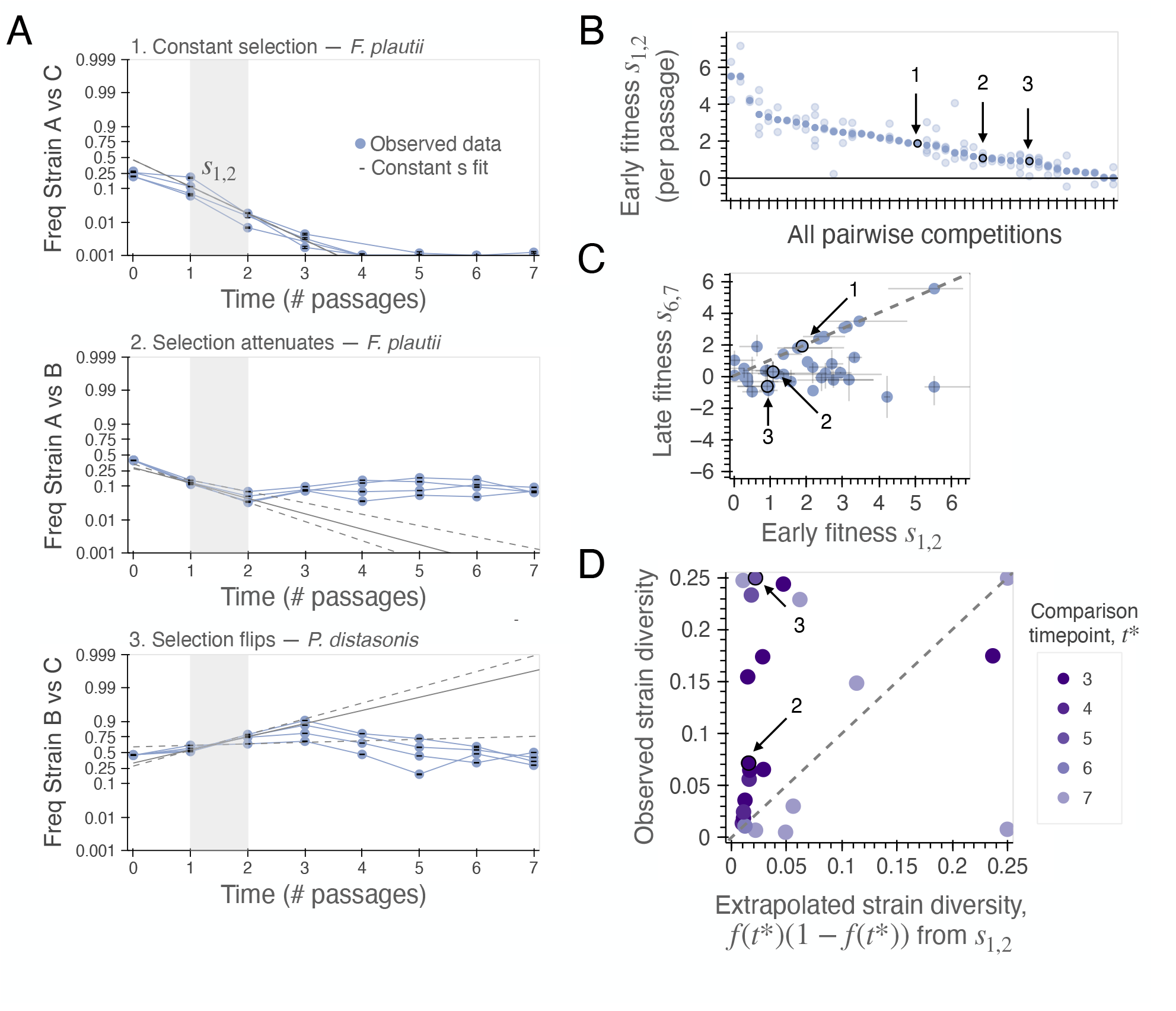
Selection pressures shift over time, promoting strain coexistence. **(A)** Examples of constant and time-varying selection within species. Points denote biological replicates, while grey lines show extrapolated trajectories using the instantaneous relative fitness inferred from timepoints 1 and 2 (solid=median, dashed=upper and lower estimates; Methods). **(B)** Estimated selection coefficients between timepoints 1 and 2 for all 42 strain competitions with sufficient sequencing coverage at timepoints 1, 2, 6 and 7. Light points show individual replicates, while darker points indicate the median. Arrows denote the three examples in panel A. **(C)** Shifts in relative fitness over time for all strain pairs that remained above the limit of detection at passage 7. Symbols denote the median selection coefficient across replicates, while grey lines represent the corresponding range. Dashed line denotes the expectation if selection were constant over time. **(D)** Temporal shifts in selection tend to favor the maintenance of strain diversity. Points denote the median strain frequency at a critical timepoint *t*^*^ where the extrapolated frequency under the constant selection model was last above 1%. The excess of datapoints above the 1:1 line indicates that the net changes in *s*(*t*) were biased in the direction of increasing intra-species diversity (p<0.05, Methods).

Some in vitro competitons were consistent with this prediction, with the fitter strain sweeping within the focal species at a roughly constant rate (e.g. *Flavinobacter plautii* in Fig. 3A). However, in many cases, we observed that the strain frequencies strongly deviated from a model where selection was constant over time. Of the 42 competitions in Fig. 3B with sufficient sequencing coverage at passages 6 and 7, more than half of the individual replicates (n=70/111, 63%) exhibited evidence for an altered selection coefficient in the second time interval (q<0.05 and |*f*_*pred*_ *-f*_*obs*_|*>*1%, Methods; Fig. S19). These temporal deviations were often consistent across multiple independent replicates (q<0.05, t-test; Figs. 3A, S20, and S21), suggesting that they were likely caused by deterministic shifts in the intra-species selection landscape over the course of the competition. Since the abiotic environment was held fixed in these experiments, we conclude that the changing intra-species selection coefficients must have been driven by shifts in the biotic environment – a multi-species analogue of frequency-dependent selection. Such feedbacks could arise through pairwise interactions between the focal strains, as well as more general interactions with other species or metabolites in the community (*79)*.

Aggregating across species, we found that the strength of selection often attenuated over time, such that the fitness differences at later timepoints were generally smaller than in the first two passages (Figs. 3C and S22; permutation test on the difference in medians, p<0.001). In some of these cases (e.g. *F. plautii* in Fig. 3A), the selection coefficients appeared to relax towards zero, in a manner suggestive of stable coexistence (*79)*. In other examples (e.g. *P. distasonis* in Fig. 3A), the direction of selection reversed at later timepoints, such that both strains remained at intermediate frequencies by the end of passage 7. Interestingly, we found that in aggregate, these shifts in relative fitness tended to be biased in the direction of increasing intra-species diversity. By comparing the number of competitions that fell above versus below their corresponding constant selection curves (Figs. 3D and S23), we found that a majority of the deviations (n=16/23) were biased towards leaving the strains at intermediate frequencies (p<0.05, permutation test, Methods; Fig. S24). This suggests that the time-varying selection coefficients tended to promote coexistence of conspecific strains within the context of their larger community, in a manner reminiscent of negative frequency-dependent selection.

### Conspecific strains exhibit fitness tradeoffs across different abiotic environments

Our experimental design also followed independent replays of the same collision across two different nutrient environments (Fig. 4A). This allows us to ask how the dynamics of intra-species competition shift under changes in the abiotic environment. Previous theoretical work has suggested that large communities might self-organize to shield themselves from external environmental perturbations (*40, 84, 85)*, but the extent to which this occurs in practice remains unclear.

**Fig. 4:**
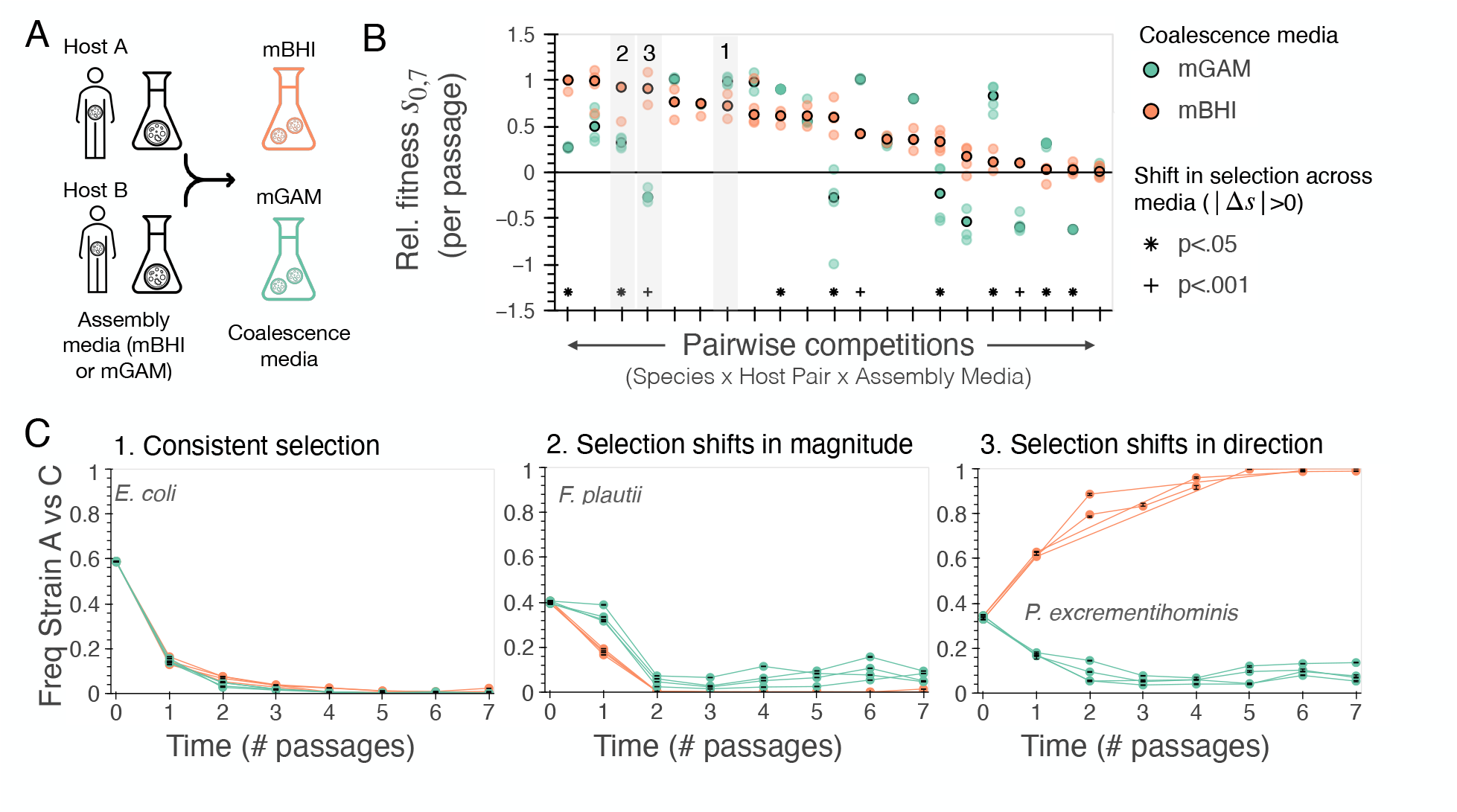
Conspecific strains exhibit fitness tradeoffs across different abiotic environments. **(A)** Community coalescence experiments were performed in two different nutrient conditions, allowing comparisons of strain-level fitness across different abiotic environments. **(B)** Net selection coefficients between passages 0 and 7 for all pairwise competitions that could be measured in collisions from both media. Dark and light points show medians and individual replicates as in Fig. 3D. Competitions with significant differences between environments are indicated (FDR corrected t-test on frequencies at passage 7). **(C)** Examples of consistent and variable selection pressures across the two abiotic environments. All three examples were obtained from the collision between hosts A and C.

To investigate this question, we focused on the 21 cases where the same species was present at sufficient abundance in collisions performed in both growth media. We then asked how the underlying strain trajectories in these species differed from each other. In some cases, such as the *E. coli* competition in Fig. 4C, we observed remarkably consistent trajectories across the two abiotic environments, both in the qualitative shape of the trajectory as well as the quantitative values of *s*(*t*). In other cases, we observed significant differences in the strain trajectories across the two abiotic environments. Some of these shifts were gradual in nature: for instance, the *F. plautii* strain from host A declined in frequency in both media when competing against the strain from host C (Fig. 4C), but the magnitude of the decline was greater in mBHI than in mGAM (q<0.05, t-test). In other cases, the direction of selection flipped between the two abiotic conditions (e.g. *P. excrementihominis* in Fig. 4C, q<0.001, t-test), or disrupted instances of apparent coexistence (e.g. *P. excrementihominis* in Fig. S25, q<0.001, t-test).

To quantify the prevalence of these effects across the broader set of strain competitions, we computed the net selection coefficients between passages 0 and 7 and compared them across the two nutrient environments (Fig. 4B; Methods). We observed significant environmental differences in the net selection coefficients in ∼50% of cases (n=11/21; q<0.05, t-test), half of which (n=5/11) involved a reversal in the overall sign of selection (Fig. S26). Interestingly, these fitness differences were not strongly correlated with the initial relative abundances of the strains (Fig. S27), or their shift in relative abundance across the two conditions (Fig. S28). These data show that intra-species selection can be highly contingent on shifts in the abiotic nutrient environment. In a community context, these genotype-by-environment interactions could be mediated both by direct effects of the growth media on the focal strains, or by indirect effects resulting from shifts in the abundances of other community members (Fig. S10).

### Deviations from transitivity reveal community-specific selection pressures

We also asked how intra-species competition depends on the biotic environment. While the results in Fig. 3 show that selection is often sensitive to the surrounding community, these shifting selection coefficients could be caused by biotic interactions that are common to all communities assembled in the same environment, as well as by idiosyncratic interactions (e.g. the identity of the opposing strain) that vary across communities from different hosts. One can test for these latter effects indirectly by examining the transitivity of strain-level selection across collisions between different hosts. If the underlying fitness landscape was consistent across communities, then we would expect that the selection coefficients between different pairs of conspecific strains should be transitive at each point in time (Fig. 5A; 86). By contrast, any signatures of non-transitivity would indicate that the relative growth rates of these strains varied in a community-specific manner.

**Fig. 5:**
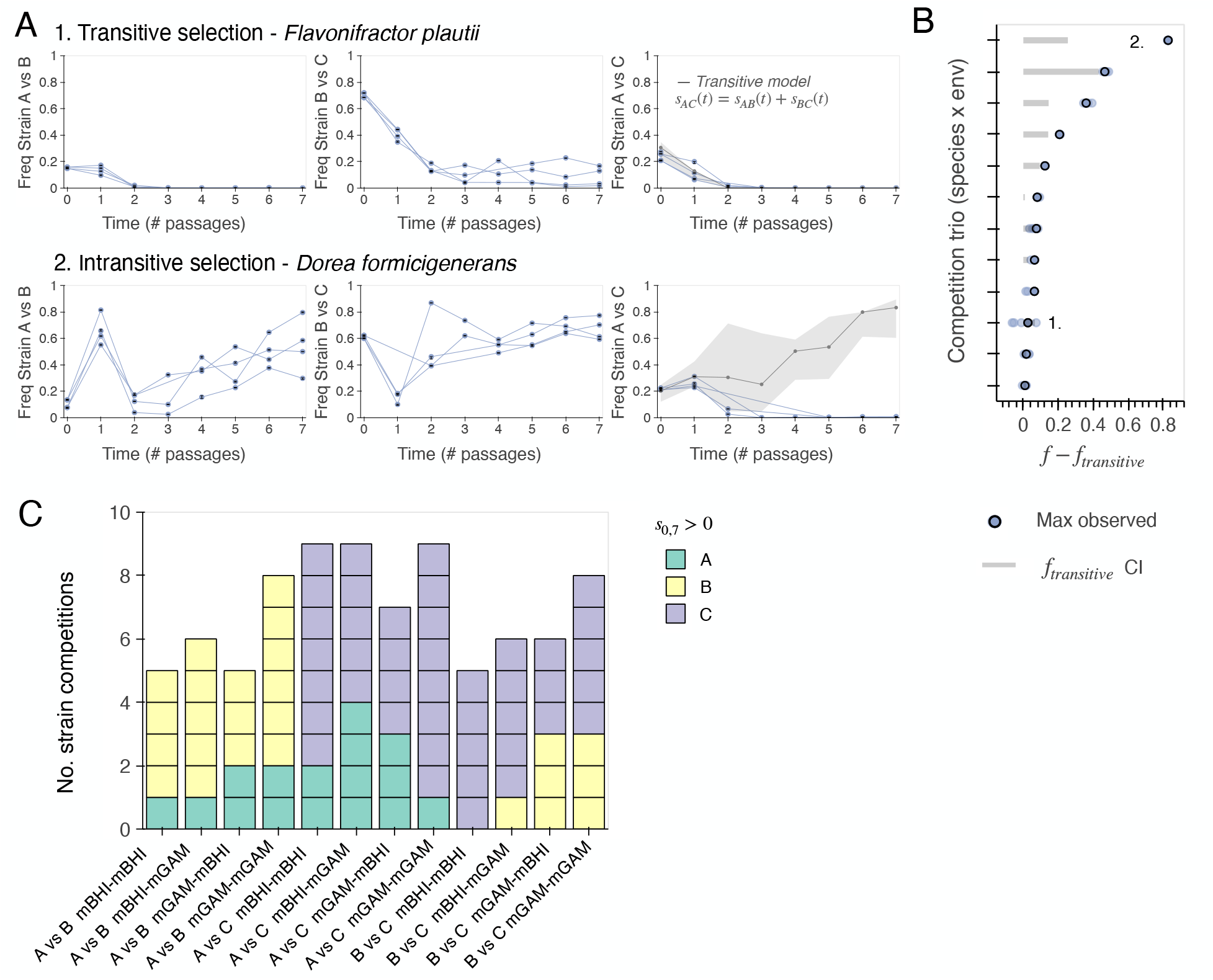
Evaluating the role of community-specific interactions on intra-species competition. **(A)** Transitivity of intra-species competition across different community collisions. Left panels show examples of strains exhibiting transitive (1) and intransitive (2) selection. Each column shows the strain trajectories for the three possible pairwise collisions. The grey lines in the third column show the predicted dynamics under a model of transitive fitness using the observed strain trajectories in the first two columns (Methods). Shaded regions represent ranges of values obtained through 100 bootstrap replicates. **(B)** Deviations from transitivity across all other species involved in a competition trio. Dark points indicate the maximum deviation between the transitive model and the median strain trajectory in panel A, while lighter points show individual replicates at the same timepoint. Grey bars show the confidence interval of the transitive model shown in panel A. **(C)** Community-wide correlations in strain-level competition. Rows illustrate different pairwise coalescence experiments, while entries are colored by the host in which the median net selection coefficient was positive. These data show that most collisions exhibited a mixture of winning strains from both hosts.

A striking example of non-transitivity is illustrated by *Dorea formicigenerans* in Fig. 5A. In this species, the competitions between hosts A-vs-B and B-vs-C suggest that the strain from host A should dominate over the host C strain if the underlying fitness landscape was transitive. However, the actual data showed the opposite trend, with the strain from host C increasing in frequency over the host A strain in a rock-paper-scissors-like manner (*s*_*AB*_, *s*_*BC*_ *>* 0 but *s*_*AC*_ *<* 0; Fig. 5A).

These rock-paper-scissors-like dynamics were uncommon among the broader collection of species in our dataset. Of the 12 cases where all three pairwise competitions could be observed, the *D. formicigenerans* example in Fig. 5A was the only one where the sign of selection clearly flipped. We therefore tested for more quantitative deviations from transitivity by leveraging the relative fitness model in Eq. (1). For each of the 12 trios, we used two pairwise competitions to infer the relative fitness trajectories for two of the three pairs of strains at each point in time (Methods). Under the assumption of transitivity, this yields a time-varying prediction for the third pairwise competition (shaded region in Fig. 5A) that has only one additional parameter (corresponding to the initial relative frequency). By comparing these predictions against the data (Fig. 5B), we identified some cases (n=6/12) where the observed trajectories closely matched the predictions of the transitive model (Figs. 5A and Fig. S29). However, we also observed several additional cases (n=6) in which there were quantitative deviations from transitivity even when the overall sign of selection was preserved (Figs. 5A and S30; Methods). These results, together with our earlier observations in Figs. 2D and 3, demonstrate that strain-level competition can be contingent on the local biotic context, even in communitiesoriginally assembled in the same abiotic environment.

### Limited strain-level cohesion during community coalescence

Finally, we sought to examine whether strain-level competition was correlated across multiple species in the same community. Such correlated outcomes could arise from direct interactions between strains, e.g. if one strain consumes a metabolite produced by another (*73, 75)*. They could also arise indirectly if the strains one community are better adapted to the environment where community coalescence occurs (*77)*.

To test for these community-wide correlations, we calculated the net selection coefficient between timepoints 0 and 7 for all strains in a given collision, and asked whether the sign of selection was biased towards one subject of origin or the other (Figs. 5C and S31; Methods). We found that the winning strains in each collision often originated from different hosts, such that the average host bias was relatively small (IQR of binomial probability, 0.59-0.83). While no individual collisions were significant after correcting for multiple comparisons (q>0.05, permutation test of G-statistic; Methods), we observed a small but significant shift in the median bias compared to a null model where the winners were randomly chosen from each host (p=0.002; Methods, Fig. S32). This suggests that strain-level cohesion played a limited role in our experiments, despite their history within each host and their co-selection during the initial round of in vitro community assembly.

## Discussion

Our results show that community coalescence can provide a scalable approach for measuring intra-species competition within larger microbial communities. By colliding in vitro gut communities derived from healthy human donors, we quantified the emergent selection pressures on conspecific strains across a range of biotic and abiotic conditions. Our statistical characterization of this landscape revealed that strains from different hosts are generally subject to strong and context-dependent selection pressures, even when their parent communities were originally assembled in the same abiotic conditions. We also found that the relative fitnesses shifted over time due to ecological interactions within the community, enabling long-term coexistence of multiple conspecific strains in the same spatial location. While these observations do not directly establish the underlying mechanisms of coexistence, they nevertheless support the view that some conspecific strains can coexist with each other through local niche partitioning.

Our results show that this intra-species coexistence can occur in large communities that appear to be “ecologically saturated” at the species level, with the total number of surviving species remaining close to the initial pre-collision baseline (Figs. 2B and S12). However, we also found that conspecific strains generally experience strong selection, even when the relative abundance of their focal species is comparatively stable (Figs. 2D,E and S16). Further work is necessary to determine whether these species- and strain-level selection pressures are governed by similar ecological mechanisms (e.g. resource competition; 41, 53, 87, 88), and how they extend to the lower-abundance taxa that are missed by our current strain-tracking pipeline. In either case, our results suggest alternative strategies for engineering targeted community engraftment events, by leveraging the existing phenotypic diversity across large panels of conspecific strains from different hosts.

Our results are consistent with recent theoretical predictions showing that mutant strains can often coexist within larger communities without exclusive spatial or metabolic niches (*40, 89)*. However, on longer evolutionary timescales, the conspecific strains in our study exhibit few genome-wide signatures of stable ecological differentiation (or “ecotypes”) (*90)*, with high levels of gene flow that are more consistent with a single biological species (*20, 21, 23, 47)*. Understanding how this long-term genetic cohesion emerges against a backdrop of widespread local niche differentiation remains an interesting evolutionary puzzle. Our results suggest that local coexistence could help facilitate this genetic mixing in ecologically diverse communities like the human gut, by providing more opportunities for conspecific strains to engage in horizontal gene transfer and recombination (*21, 55, 91)*.

Our competition experiments were performed in a simplified environment that omits important factors, such as spatial structure or host immunity, that are thought to influence strain-level selection in natural gut microbiomes. While these simplifications allowed us to show that conspecific strains can coexist even in the absence of these additional factors, further work is needed to understand how the fitness differences measured in our community experiments relate to the specific selection pressures that these bacterial strains experience in vivo. Future experiments could start to incorporate these factors in a controlled manner, by performing similar community collisions in spatially structured environments (*71)* or in gnotobiotic mice (*92)*.

Our experiments also focused on the standing genetic variation present among gut bacterial strains from different hosts. On longer timescales, these initial strains will start to acquire further genetic changes as they begin to adapt to their local community environment (*12, 82, 83)*. These processes of local evolution and inter-strain competition, when aggregated over the larger collection of human microbiomes, ultimately generate the phenotypic and genotypic diversity within our initial strain pool. Understanding the evolutionary rules that shape this phenotypic landscape —and how they impact competition on local scales — will be critical for understanding how large microbial communities evolve.

## Materials and methods

### Collection of stool samples

This study was approved by the Stanford University Institutional Review Board as Protocol IRB-64602, and written, informed consent was obtained from all participants. Healthy adult individuals were recruited from Stanford University and the surrounding community. Exclusion criteria included age under 18, pregnancy or nursing, chronic illness and routine use of prescription medications known to influence the gastrointestinal tract, as well as antibiotic usage in the past three months. Stool samples were collected using the protocol described in Ref. (*25)*. Each participant collected one stool sample by depositing ∼20 mL of stool into a vial. Stool samples were frozen immediately after collection in donors’ home freezers and were stored there for up to one week until they were transferred to storage at -80 °C in the laboratory. We initially collected stool samples from 8 study participants (Tables S1 and S2). After an initial round of sequencing and species profiling (see below), we selected 4 of these samples for further in vitro passaging based on the abundances of their *Bacteroidaceae* species.

### Assembly of in vitro communities from stool samples

In vitro communities were assembled from frozen stool samples using the methods described in Refs. (*41, 65, 66)*. All culturing experiments were performed in an anaerobic chamber (Coy instruments, H_2_ 2.4%, CO_2_ 6.5%) with pre-reduced media and solutions. Small fragments of frozen stool (∼ 50 mg) were first resuspended in 1 mL of pre-reduced, filter-sterilized PBS. This initial stool suspension was then used to create six initial cultures, by adding 20 *µ*L aliquots of the stool suspension to 180 mL of pre-reduced mGAM (HyServe No 1005433) or Brain-Heart Infusion (BHI, BD Biosciences 237200) supplemented with 0.2 mg/mL L-tryptophan, 1 mg/mL L-arginine, 0.5 mg/mL L-cysteine, 5 *µ*g/mL vitamin K, .5 *µ*g/mL hemin (mBHI) in a 96-well deep-well plate, for a total of three replicates per subject per media condition. These two media were chosen because they had previously been shown to support the growth of taxonomically diverse communities (*65–68)*, while differing in the specific taxa that are favored in each condition. After inoculation, communities were grown at 37 °C and passaged via a 1:200 dilution into fresh pre-reduced media every 48 hr for a total of 5 passages. In the absence of cell death, this corresponds to an average of ∼40 generations of total growth, assuming log_2_(200) ≈7.7 cell doublings per passage; the actual number will vary across taxa depending on their relative growth within each cycle. Cultures were not shaken during growth, but were mixed via pipetting immediately prior to each dilution. At the end of the experiment, 50 *µ*L of each saturated culture was frozen in 25% glycerol and stored in mylar bags at -80 °C for future use. This yielded a total of 24 initial communities (Table S3).

### Community coalescence experiments

Coalescence experiments were performed using a modified version of the protocol described in Ref. (*41)*. For each nutrient condition, thawed glycerol stocks from one replicate of each community at passage 5 were inoculated into 200 *µ*L of fresh, pre-reduced media and were passaged as before for two additional growth cycles. In the second growth cycle, communities were grown in a larger volume (475 *µ*L) to produce sufficient cell counts for the coalescence experiment. At the end of the second growth cycle, pairwise mixtures were formed by mixing 100 *µ*L of each saturated culture to create an initial inoculumn, with two replicate inocula created for each pair of “parent” communities. From each inocolumn, four replicate coalescence experiments were founded by adding 1 *µ*L of the initial inoculumn to 199 *µ*L of fresh mGAM (two replicates) or mBHI (two replicates); the rest of the inoculumn was frozen at -80 °C for sequencing. We also founded single-community control lines (two replicates per growth medium) using 1*µ*L of the saturated culture at the end of the second growth cycle. This yielded a total of 96 collision experiments and 32 controls (see Table S4). Communities were then propagated for seven additional passages using the protocol described above, with frozen samples preserved at the end of each passage for sequencing.

### DNA extraction and metagenomic sequencing

For the initial sequencing of stool samples, DNA was extracted from a small fragment of stool (∼50mg) using the Qiagen DNAeasy PowerSoil Kit. This same extraction method was also used to obtain DNA from the resuspended stool inocula that were used to found our initial in vitro communities. DNA from subsequent passages was extracted from 50 *µ*L of frozen culture using the Qiagen DNAeasy Ultraclean 96 Microbial Kit. Metagenomic sequencing libraries were prepared using the Nextera DNA Flex Library Prep kit. Sequencing was performed on an Illumina NovaSeq with read lengths of 2×150bp and a target depth of ∼5 Gbp per sample (Table S1). This target depth was chosen so that species with a relative abundance of >1% would have an expected coverage of >5x, enablng measurements of their intra-species diversity (see below).

### Species and SNP profiling of metagenomic samples

Raw sequencing reads were trimmed using Skewer (*93)* and reads mapping to the human genome (GRCh38) were filtered using Bowtie 2 (*94)*. Remaining reads will be made available in the SRA upon publication.

Species relative abundances were estimated using the species module of MIDAS2 (*95)*, which maps the trimmed sequencing reads against a panel of universal, single copy marker genes representing different bacterial species from the UHGG database (*5)*. Relative abundances were reported for all species that had at least two marker genes with at least two mapped reads in a given sample. We used these estimated species abundances for all species-level analyses, such as calculating species richness or Jensen-Shannon distance (see below).

Single nucleotide variants (SNVs) were identified using a variant of the MIDAS2 snps module. We first generated a list of the 150 most abundant species, based on their maximum relative abundance across all samples. We then used the MIDAS2 snps module to map the raw sequencing reads from each sample against a set of dereplicated reference genomes representing these different microbial species. For each site in the associated reference genomes, MIDAS2 reports the number of reads mapping to each allele, as inferred from the read pileups in that sample. To enable accurate tracking of mutations over time, we modified the source code of MIDAS2 to report read counts for all alleles, independent of their total count within a single sample. We then used the MIDAS2 merge module to tabulate allele counts across all samples in our study. We only included populations with a median vertical depth of 1 and a horizontal depth of 0.5, and excluded all sites whose total coverage differed from the median vertical depth by more than 2.5-fold. For our downstream analyses, we restricted our attention to populations where the median non-zero coverage along the genome 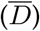 was greater than 5x. At our targeted sequencing depth of ∼5 Gbp per sample, this translates to a minimum relative abundance of ∼1%. Our strain-level analyses were therefore restricted to those taxa that were consistently present above this threshold (Table S5). Further work is necessary to determine how these results extend to rarer taxa that are not able to be profiled with this method.

### Quantifying species composition and community stability

For each sample, we estimated the total species richness using the number of species that were present with an estimated relative abundance greater than or equal to 0.1%. We also computed the compositional dissimilarity between pairs of communities using the Jensen-Shannon distance (JSD),

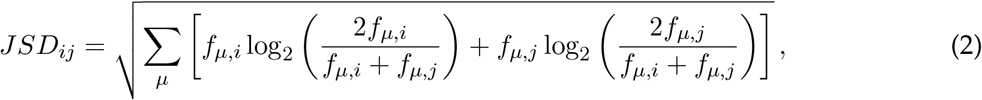

where *f*_*µ,i*_ is the relative abundance of species *µ* in host *i*. We only used samples in which at least 1000 reads mapped to the database of species marker genes.

We used these metrics to quantify the time to stabilization during community assembly and coalescence (Figs. 1BC, 2BC, S2A, S8, and S9). To assess convergence using species richness, we compared the distribution of species richness at a given timepoint to the distribution at the final timepoint (passage 5 for community assembly and passage 7 for community coalescence). Statistical significance was assessed via permutation tests of the median values of each distribution.

To assess convergence using JSD, we used a similar procedure to compare the distribution of JSDs between the two final timepoints (4-5 for assembly, and 6-7 for coalescence) to the distribution of JSDs between pairs of earlier consecutive timepoints. We also quantified community specificity by comparing the distribution of JSDs between pairs of replicate communities to the JSDs of communities assembled or coalesced in different conditions.

### Quantifying intra-species diversity within and between samples

For each species population in each sample, we estimated the intra-sample genetic diversity by calculating the fraction of non-filtered sites in which the major allele was present at an intermediate frequency (*f <* 0.8). We also calculated a measure of the genetic divergence between pairs of samples, defined as the fraction of non-filtered sites in which an allele was present at high frequency (*f >* 0.8) in one sample and low frequency (*f <* 0.2) in the other (Fig. S13A). Many in vitro populations had an intra-sample diversity much lower than their average divergence to samples from other hosts (Figs. 1D, 1E, S3A, and S13A), suggesting that these samples were likely dominated by a single colonizing strain (*9, 69)*. Following Ref. (*9)*, we used these data to classify a given in vitro population as effectively mono-colonized (“quasi-phaseable”) if its intra-sample diversity was less than 10^*-*3^, which is approximately 10-fold lower than the typical between-subject divergence in Fig. 1E. This threshold was designed to detect cases where two (or more) typically diverged strains are present at frequencies greater than ∼10% (*9)*.

### Testing for cross-contamination between samples

We checked for signatures of cross-contamination during passaging by maintaining a set of blank wells on each 96-well plate. At each passage blank wells were visually inspected for microbial growth to confirm that no contamination had occurred. We also checked for signatures of contamination in our genetic data using generalization of the approach in Ref. (*25)*, which searches for unexpected instances of strain sharing between communities that were derived from different hosts. Our passaging experiments were organized so that many neighboring wells contained communities that were assembled from different hosts, thereby increasing the chances of detecting cross-contamination that may have arisen during passaging or sequencing. Strain sharing between communities was assessed by computing the genetic divergence metric defined above for each species, and searching for cases in which we observed fewer than 10 fixed differences (sites where an allele is present at >80% frequency in one sample and <20% frequency in another). Although it is possible that unrelated subjects can contain genetically similar strains by chance, previous work has shown that two random subjects are unlikely to share multiple highly similar strains (*9, 25)*. We considered there to be sufficient evidence for contamination if multiple species showed evidence of strain sharing with a sample from another subject (excluding the collided subject in samples from the community coalescence arm). This procedure revealed that one of the four initial fecal suspensions was likely contaminated with bacteria from another subject. To be conservative, we removed all samples involving this subject from all of our downstream analyses. This left a total of three initial hosts, with 48 community coalescence experiments and 24 autologous controls.

Since fixed differences metrics can be less sensitive for detecting strain sharing between samples with multiple conspecific strains (*9, 25)*, we also used an allele sharing metric (*70)* to confirm that the instances of multi-colonization in Fig. 1D were not driven by contamination. For each shared species in each pair of samples, we calculated the number of sites in which the non-reference allele was present above 20% frequency in both samples (Fig. S3B). Following Ref. (*25)*, we considered there to be sufficient evidence for contamination if multiple species showed high levels of non-reference SNV sharing (>75 % SNVs shared).

### Tracking the relative frequencies of conspecific strains during community coalescence

For each community coalescence experiment, we identified the subset of species that were present at sufficient coverage (*D >* 5) in both parent communities immediately prior to mixing. We then quantified the relative frequencies of the strains within these shared species during the resulting coalescence experiment. Strain frequencies were estimated using a generalization of the “quasi-phasing” approach in Refs. (*9)* and (*25)*, leveraging the fact that both parent communities were measured separately prior to mixing.

For most of our analyses, we restricted our attention to species where both parent populations were classified as quasi-phaseable according to the criterion described in the previous section (i.e. within-sample diversity *<* 10^*-*3^). For these populations, previous work has shown that the genotype of the dominant strain can be inferred with a high degree of confidence, by searching for alleles that are present at a sufficiently high frequency (e.g. *f >* 0.8; 9). We used this idea to identify a collection of marker SNVs that distinguish the dominant strains in each of the pre-mixture communities. A SNV was counted as a marker SNV for a given strain if the non-reference allele was present at >80% frequency in its pre-mixture population and <20% frequency in the other population that it was mixed with. We further refined these sets of marker SNVs by removing any SNVs that deviated from the median marker SNV frequency by >50% frequency in >25% of the samples from the coalescence experiment; Fig. S13B shows that this typically leaves ∼ 1000 putative marker SNVs from each pairwise collision. The resulting sets of marker SNVs were then used to track the relative frequencies of the conspecific strains during coalescence.

The frequency of each strain was estimated in each subsequent timepoint using a piecewise estimator designed to reduce the impact of mapping errors and other sequencing artifacts. We first computed the median frequency across all the the marker SNVs for a given strain,

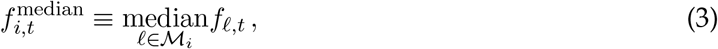

where *f*_*l,t*_ is the frequency of the non-reference allele at site *l* in timepoint *t* and ℳ _*i*_ is the set of marker SNVs corresponding to strain *i*. We then estimated the frequency of strain *i* using the piecewise formula,

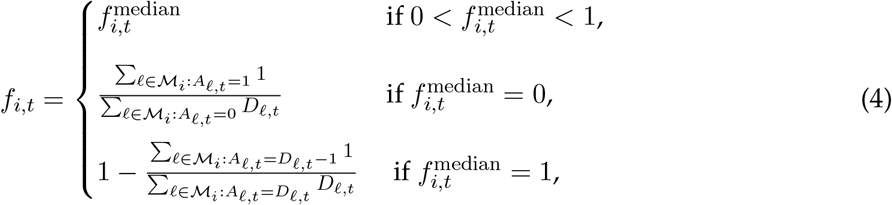

where *A*_*l,t*_ is the number of reads supporting the alternate allele at site *l* at timepoint *t* and *D*_*l,t*_ is the total coverage. In other words, Eq. (4) estimates the frequency of a strain as the median frequency of its set of distinguishing snps, unless the median frequency was 0 or 1; in those cases we estimated the frequency of a strain using the ratio of counts of sites with 0 reads vs 1 read of the appropriate allele. We then re-normalized the relative frequencies of strains such that *f*_*i,t*_+*f*_*j,t*_ = 1. Deviations from this normalization can be caused by sequencing noise as well as the presence of other low frequency strains. To be conservative, we eliminated all timepoints where *f*_*i,t*_ + *f*_*j,t*_ deviated from the expected normalization by more than 10% (Fig. S13C), which is larger than what would be expected from sampling noise alone.

We assessed the uncertainty in our frequency estimates by bootstrapping sets of marker SNVs. We generated 1000 bootstrapped replicates for each strain frequency and estimated the 95% confidence intervals using the 2.5th and 97.5th percentiles of the bootstrapped distribution. We used these estimates to plot the confidence intervals in Fig. 2E.

### Validation on simulated metagenomes

To evaluate the limits of our strain-tracking approach, we applied our frequency estimation pipeline to simulated metagenomic datasets designed to mimic the conditions of our experiment. For each simulated metagenome, we used the read simulator InSilicoSeq v2.0.1 (*96)* to generate ∼7 million paired end reads under the NovaSeq error model. Simulated metagenomes were constructed using genomes from the UHGG database (*5)* using the observed species abundances from two different coalescence experiments, both of which contained a population of *Bacteroides thetaiotaomicron* (*Bt*). For all species except *Bt*, we generated synthetic reads from the corresponding MIDAS2 reference genome (*95)* in proportion to their observed frequency in the sample. For the *Bt* populations, we simulated multi-strain population by generating a mixture of reads from the *Bt* reference genome and another *Bt* genome from the UHGG database (GUT_GENOME000472 or GUT_ GENOME001637), which were chosen to span a range of genetic distances from the reference. We simulated a range of mixture ratios from 1:10^5^ to 10^5^:1 while varying the coverage of the *Bt* population from 5x-30x (by altering the relative abundance of *Bt* and re-normalizing the abundances of the other species). We then estimated the frequencies of the non-reference *Bt* strain using the procedure described above (Fig. S14).

As expected, the accuracy of the frequency estimates strongly depended on the corresponding number of marker SNVs. For the more diverged *Bt* strain, we typically identified around 20,000 marker SNVs, and could accurately detect the strain at frequencies as low as 10^*-*5^ when the Bt coverage was at least 5x. By contrast, the less-diverged *Bt* strain typically yielded around 200 marker SNVs, and the resolution of our frequency estimates plateaued around ∼10^*-*3^ for the lowest depth samples. We used these simulation results to establish the thresholds for our frequency estimation pipeline. In particular, we required each species to have a total coverage of at least 5x to perform strain frequency inference, and set our strain detection floor to *f*_min_ ≡10^*-*3^.

### Quantifying strain-level selection coefficients

We used the frequency estimates above to compute the relative fitness between pairs of strains using the formula in Eq. (1). For a given pair of timepoints *t*_1_ and *t*_2_, we estimated the average selection coefficient within that window using the formula

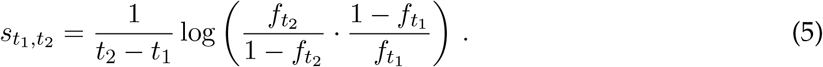

Since this equation is undefined for *f*_*t*_ = 0 or *f*_*t*_ = 1, we capped the strain frequencies at our estimated detection limit of *f*_min_ *≡* 10^*-*3^ to avoid arithmetic overflow.

We used these estimates to test for time-varying selection by examining whether selection an an earlier epoch (*t*_*early*,1_, *t*_*early*,2_) was predictive of dynamics at later timepoints (*t*_*late*,1_, *t*_*late*,2_). To minimize statistical artifacts such as regression to the mean, we used a bootstrapping approach to assess statistical significance by dividing the corresponding marker SNVs into several independent groups.

For each bootstrap replicate *b*, we randomly assigned 1/4 of the marker SNVs to each of the four timepoints (*t*_*early*,1_, *t*_*early*,1_, *t*_*late*,1_, *t*_*late*,2_) by sampling without replacement, and computed a set of four independent frequency estimates, 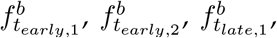 and 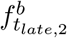. We then used these independent frequency estimates to compute an estimated selection coefficient for both epochs, which were used to plot the data points in Fig. 3C.

To test whether the shifts in *s*(*t*) tended to be biased in the direction of intra-species coexistence, we used the estimated selection coefficient from the first epoch to estimate an extrapolated frequency at later timepoints under a null model of constant selection:

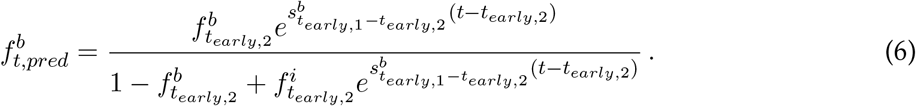

We used this extrapolated frequency trajectory to find the latest timepoint (*t*^*^) where the expected strain frequency was above a minimum threshold (*f* ^*\**^) where it could still be reliably observed. We set this threshold to *f \** = 10^*-*2^, so that it was about 10-fold higher than our estimated limit of detection from our *in silico* simulations (*f*_min_ ∼ 10^*-*3^). We then asked how the observed intra-species diversity at time *t\**, as quantified by the heterozygosity 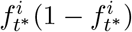 , compared to the expected diversity under the constant selection model in Eq. (6), as illustrated in Fig. 3D. Points above the 1:1 line indicate that the changes in *s*(*t*) led to a higher value of intra-species diversity, while points below the line imply the opposite. If *s*(*t*) varied in an unbiased way over time (e.g. due to the transient effects of the community collision), one would expect that a roughly equal number of competitions should fall above versus below this constant selection line. By contrast, an excess of points with higher diversity would signal that the changes in *s*(*t*) were biased in the direction of either stable or unstable coexistence.

Aggregating across species, we observed an excess of points above the constant selection prediction, both for individual competitions (Fig. S23) and when pooling across replicates (Fig. 3D). We assessed the significance of this trend by comparing the data to a null model in which each competition was equally likely to fall above or below its constant selection line. By simulating values from this null distribution, we found that the excess of points above the line was significantly greater than expected by chance, both when individual replicates were resampled independently (p<0.01), as well as when a single value was resampled for each pooled set of replicates (p<0.05).

Since the power of this test is expected to vary as a function of predicted strain diversity, we also performed an analogous version that stratified competitions by their predicted strain diversity (Fig. S24). To do so, we scanned through a continuous range of different diversity thresholds *d ∈* (0, 0.25), and computed a corresponding range of p-values that only included competitions with a predicted strain diversity ≤ *d* (Fig. S24A). We found that this p-value was minimized at an intermediate value of *d*, and significantly exceeded the null distribution obtained when this entire procedure was repeated for resampled versions of the data (Fig. S24B). This provides further evidence that the observed shifts in *s*(*t*) are biased in the direction of intra-species coexistence.

While competitions between pairs of mono-dominated populations provide the clearest estimates of intra-species selection, we tested for evidence of continued strain-level coexistence in the remaining multi-colonized populations by examining their overall levels of genetic diversity during the collision (Fig. S33). Of the 119 competitions involving two or more strains from the same host, we found that 106 (90%) continued to exhibit signatures of multi-colonization at the end of community coalescence. This fraction dropped to 17/58 (30%) among the initially multi-colonized species that did not have a competitor strain in the other host, which is comparable to the rates of intra-species coexistence among the mono-dominated species above (60/150; 40%). These observations suggest that strain-level coexistence is a widespread phenomenon in our experiments, and is not necessarily limited to pairs of strains with a prior history of co-occurrence.

In a subset of these cases, we were able to measure the underlying selection coefficients using a generalization of our marker SNV approach above. For these species, the distinguishing SNVs constitute markers for the dominant strain of the focal species in each initial community. We estimated the frequencies these strains from their marker SNVs using the same methods described above, and manually verified the resulting SNV trajectories to ensure that variants from the same strain remained correlated during the collision (Fig. S34). Within this smaller subset, we identified several additional examples of temporally varying selection (Fig. S35), suggesting that our observations of attenuated selection are not restricted to species that were mono-dominated prior to the collision.

### Quantifying signatures of non-transitive selection

We used a similar approach to evaluate the transitivity of strain-level selection coefficients between different pairs of strains. We let *s*_*A*|*B*_ (*t*) denote the (possibly time-varying) selection coefficient of the strain from host A when competing against strain B (Eq. 1). This time-varying selection coefficient could be driven by biotic factors that are common to all community collisions, as well as idiosyncratic features that are specific to the community context of one collision (including specific interactions between strains A and B). To distinguish between these scenarios, we can consider competitions with a third strain C, which yield an additional pair of selection coefficients, *s*_*A*|*C*_ (*t*) and *s*_*B*|*C*_ (*t*). If the relative fitness only depends on properties of the biotic environment that are common to all three collisions, then the selection coefficients must be transitive:

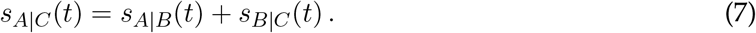

Combining this expression with Eq. (1) yields a corresponding relationship between the three relative frequencies,

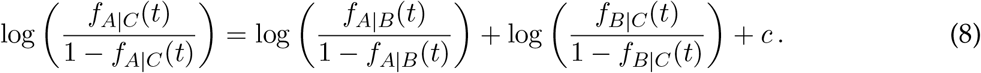

where *c* is a constant of integration that depends on the initial strain frequencies. We used this expression along with observed trajectories *f*_*A*|*B*_ (*t*), *f*_*B*|*C*_ (*t*), and *f*_*A*|*C*_ (*t*) to test for evidence of non-transitivity.

To do so, we chose the integration constant *c* to best fit the observed trajectories at timepoints 0 and 1. This yields a parameter free prediction for the remaining timepoints, which are shown in Fig. 5A. Confidence intervals for these predictions were estimated by calculating upper and lower bounds for *c* by bootstrapping replicate competitions.

### Testing for strain-level cohesion during community coalescence

We tested for evidence of strain-level community cohesion by quantifying the correlations between strain-level competition outcomes across multiple species in a given community collision. Strong correlations would correspond to scenarios where strains from a given host tend to win or lose as a group. To quantify these correlations, we calculated the net selection coefficient between timepoints 0 and 7 for all of the focal species in a given collision (Fig. 5D) and defined the winning strain as the one whose median selection coefficient across replicates was positive. We quantified the bias in these outcomes using a standard “G-statistic” metric,

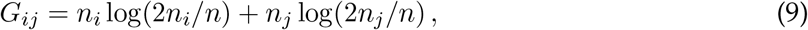

where *n*_*i*_ and *n*_*j*_ denote the numbers of winning strains from parent communities *i* and *j*, respectively, and *n* = *n*_*a*_ + *n*_*b*_ is the total number of assessed species (Fig. 5E, Fig. S32). We assessed the significance of the observed bias scores by generating bootstrapped distributions of *G* statistics by drawing *n*_*i*_ from a binomial distribution with success probability p=1/2, using the observed values of *n*. We found that the median G value across communities was not significantly elevated over the null expectation (p=0.490,Fig. S32B).

To increase statistical power, we expanded this analysis to incorporate data from initially multi-colonized populations. Repeating our cohesion analysis for this expanded set of strains yielded a small but significant elevation in the median host bias (Fig. S32A), although no individual collision was significant after correcting for multiple comparisons (q-value >0.05).

## Acknowledgments

We thank M. Bitter, L. Perez, E. Mordecai, B. Kim, T. Fukami, A. Bhatt, C. Voogdt, J.A. Lopez, and members of the Good and Petrov labs for useful discussions, and E. Visher, J. Vila, O. Ghosh, and J. McEnany for comments and feedback on the manuscript. We also thank the anonymous donors who generously contributed their fecal samples to this project. Computational analyses were performed on the Sherlock High-Performance Computing Cluster, managed by the Stanford Research Computing Center. Sequencing was performed by the Biohub – San Francisco Genomics Core.

## Author contributions

S.J.W., J.C., K.S.X., D.A.P. and B.H.G. conceived the project and designed experiments; S.J.W., R.S., H.R.G, and C.Y. performed experiments; S.J.W., Q.X., D.A.P. and B.H.G analyzed data; S.J.W., D.A.P., and B.H.G. wrote the manuscript, and S.J.W., J.C., K.S.X., D.A.P., and B.H.G. revised the manuscript with input from all the authors. All authors reviewed the manuscript before submission.

## Data and materials availability

Raw sequencing reads will be made available on the NCBI Sequencing Read Archive (SRA) upon publication at BioProject PRJNA1469869. Postprocessed data, including inferred strain frequencies and species abundances, along with scripts for data analysis and figure generation, are available on Github (https://github.com/sophiejwalton/coalescence-pilot-mgx).

## Supplementary Materials

Figs. S1-S35

Tables S1-S5

