## Supplemental Figures for "Community coalescence reveals strong selection and coexistence within species in complex microbial communities"

### Supplementary Materials

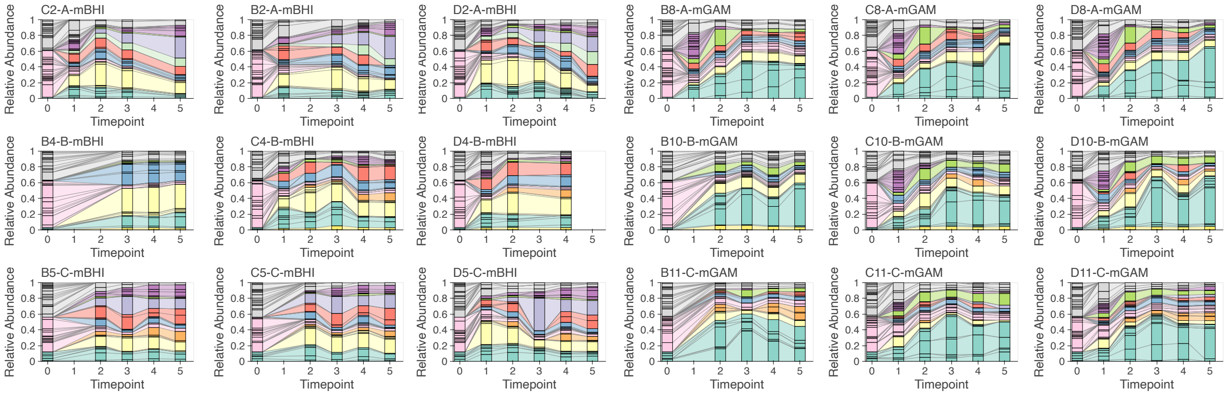


**Fig. S1: Analogous versions of Fig. 1A for all assembled communities.** Species are colored by bacterial family using the same color scheme as in Fig. 1.

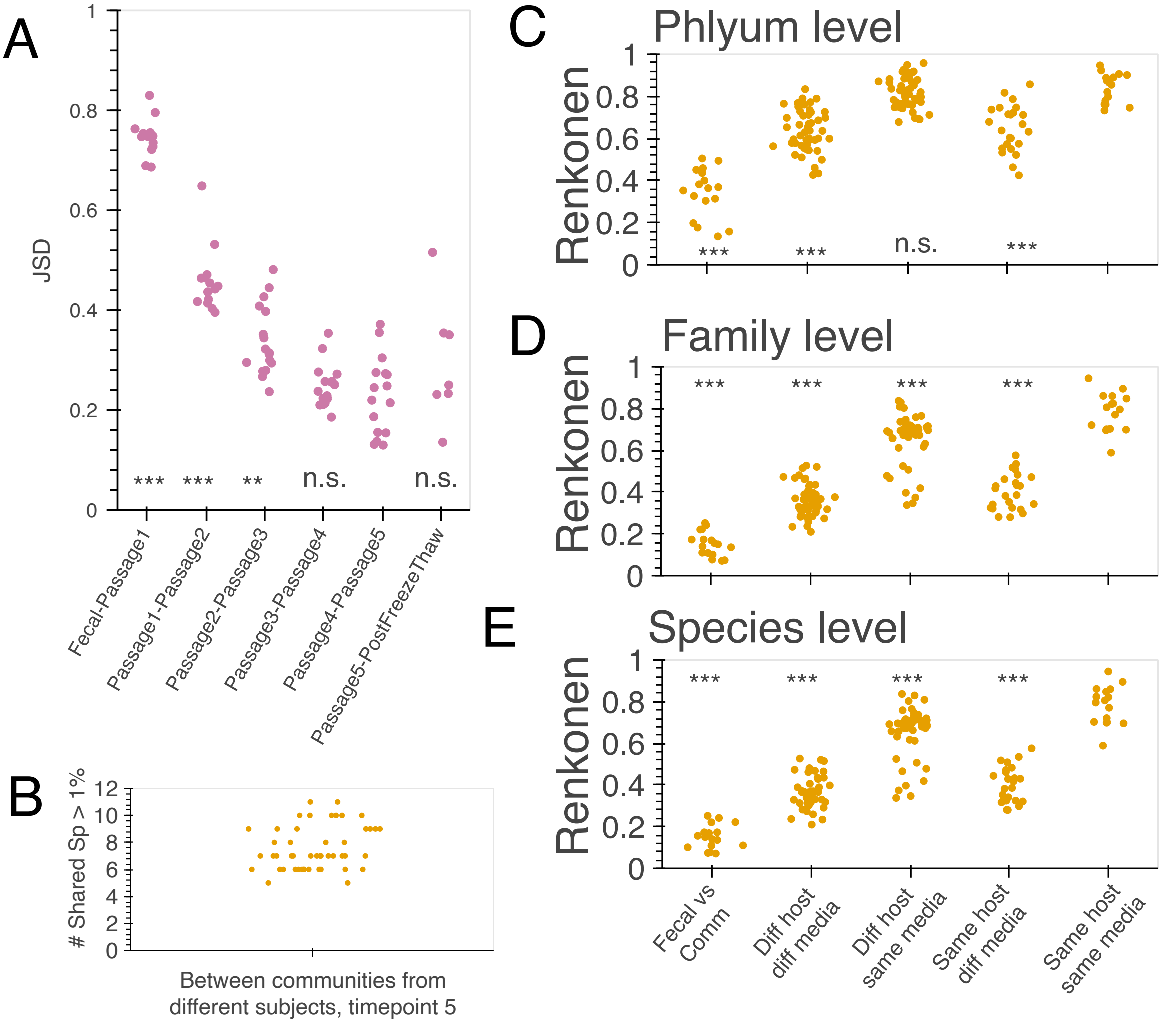


**Fig. S2: Additional characterization of in vitro community assembly. (A)** Jensen-Shannon distance (JSD) between successive timepoints of each in vitro community. Taxonomic stability was evaluated by comparing the distribution of JSD values in each column to the JSD distribution between timepoints 4-5. Asterisks show permutation tests on the difference in median JSD values (***=p<0.001, **=p<0.01, and n.s.=p>.05). **(B)** Number of species that were shared between pairs of in vitro communities assembled from different hosts with a relative abundance >1%. **(C-D)** Analogous versions of Fig. 1C using the Renkonen similarity index ($R_i j = \sum_{\mu} \min\{f_{\mu,i}, f_{\mu,j}\}$) computed for different taxonomic levels. Asterisks show permutation tests on the difference in medians compared to the biological replicates in the rightmost column (***=p<0.001).

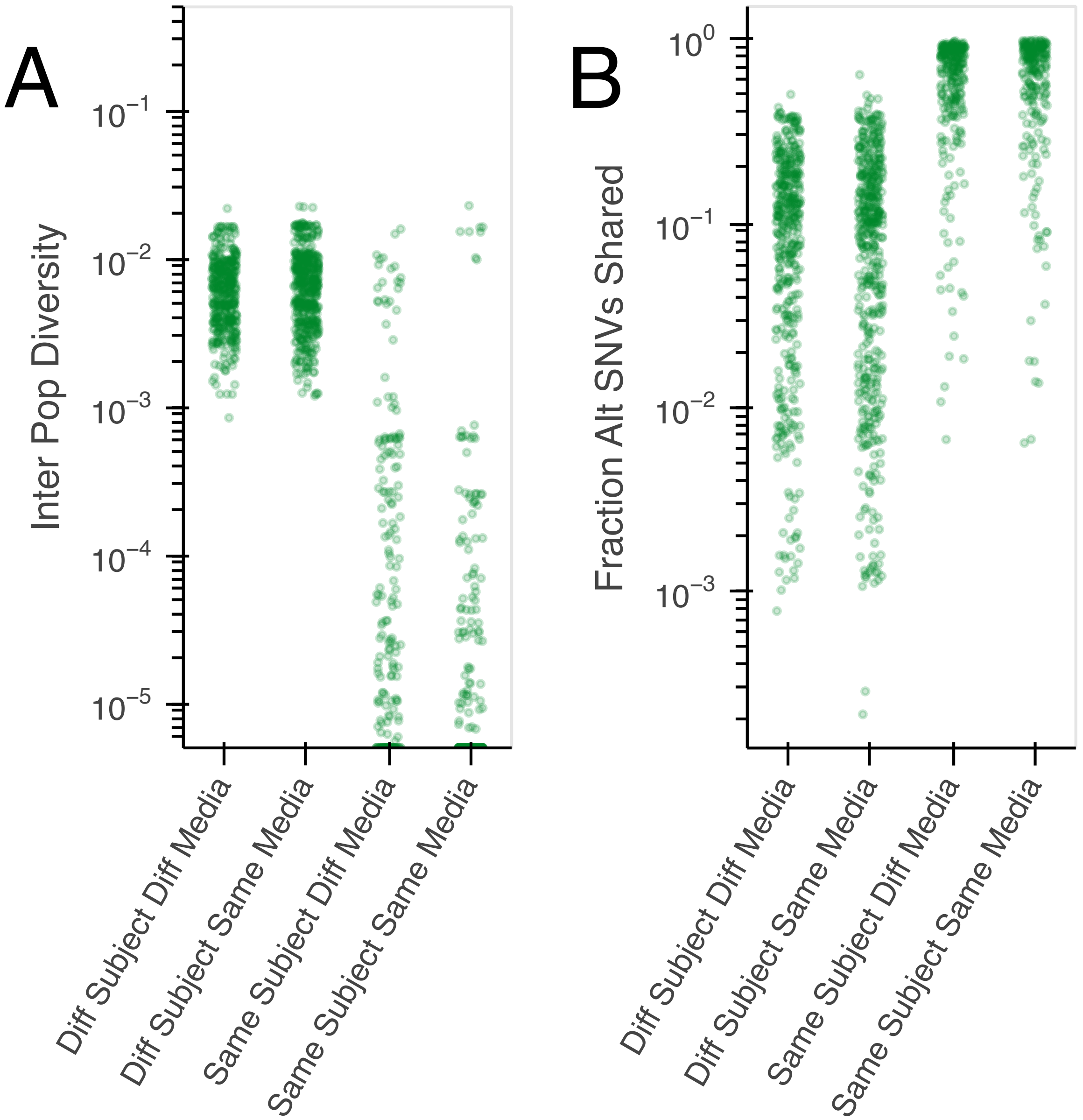


**Fig. S3: Characterization of strain sharing between in vitro communities derived from fecal samples. (A)** Analogous versions of the between-population divergence metric in Fig. 1E for different sets of in vitro populations at timepoint 5. Note that some populations from the same subject show elevated fixed differences between samples. This corresponds to cases where different strains from the initial fecal sample became the dominant strain in vitro. The outlier points in the rightmost column ('same subject, same media') include populations of *Bacteroides thetaiotaomicron* in communities assembled from host B, and populations of *Bacteroides dorei* and *Peptoniphilus harei* in communities assembled from host C. **(B)** Analogous version of panel A using a SNV sharing metric instead of fixed differences (Methods).

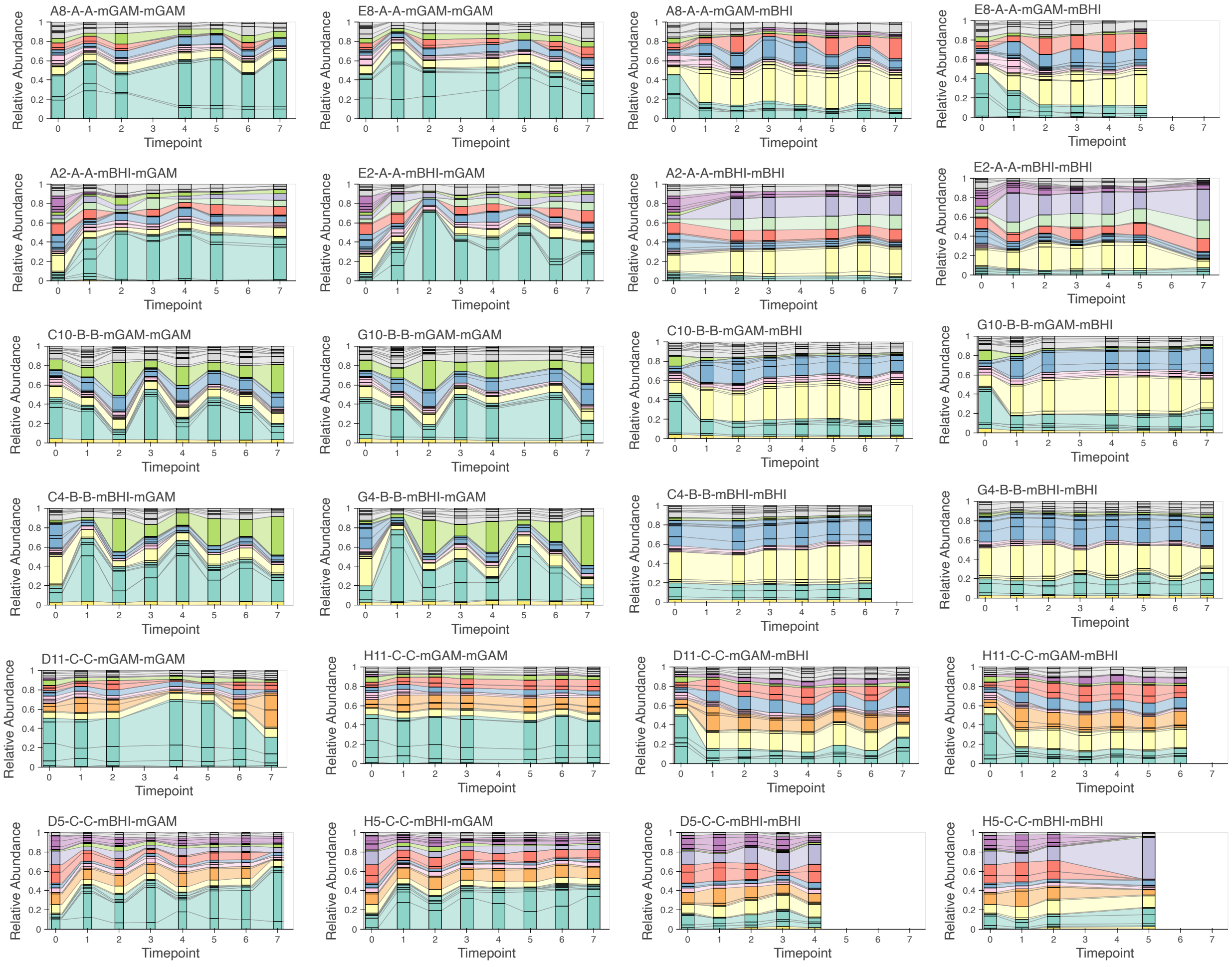


**Fig. S4: Analogous versions of Fig. 2A for the control communities in the community coalescence experiment.** Species are colored by bacterial family using the same color scheme as in Fig. 1.

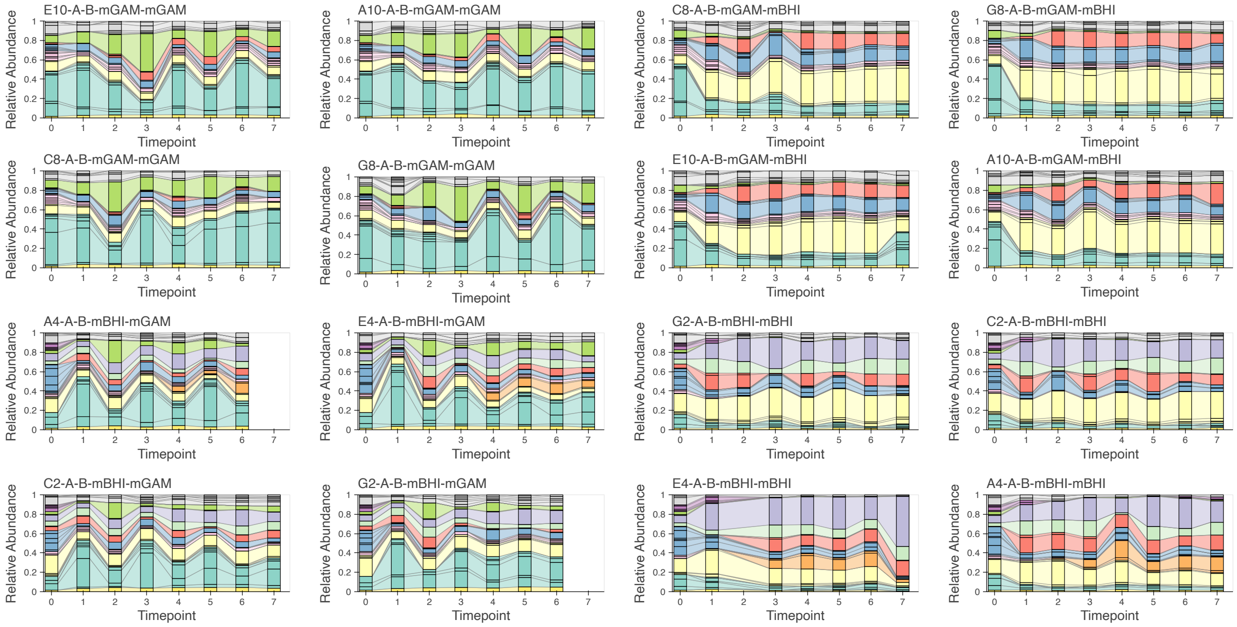


**Fig. S5: Analogous versions of Fig. 2A for all community collisions between hosts A and B.** Species are colored by bacterial family using the same color scheme as in Fig. 1.

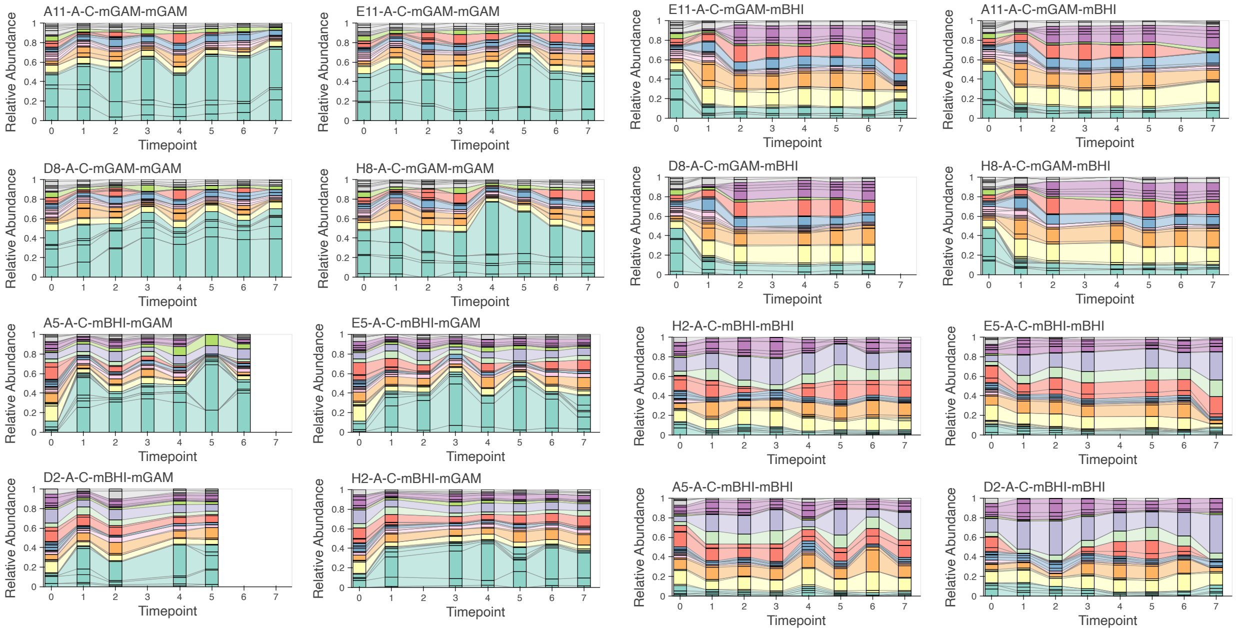


**Fig. S6: Analogous versions of Fig. 2A for all community collisions between hosts A and C.** Species are colored by bacterial family using the same color scheme as in Fig. 1.

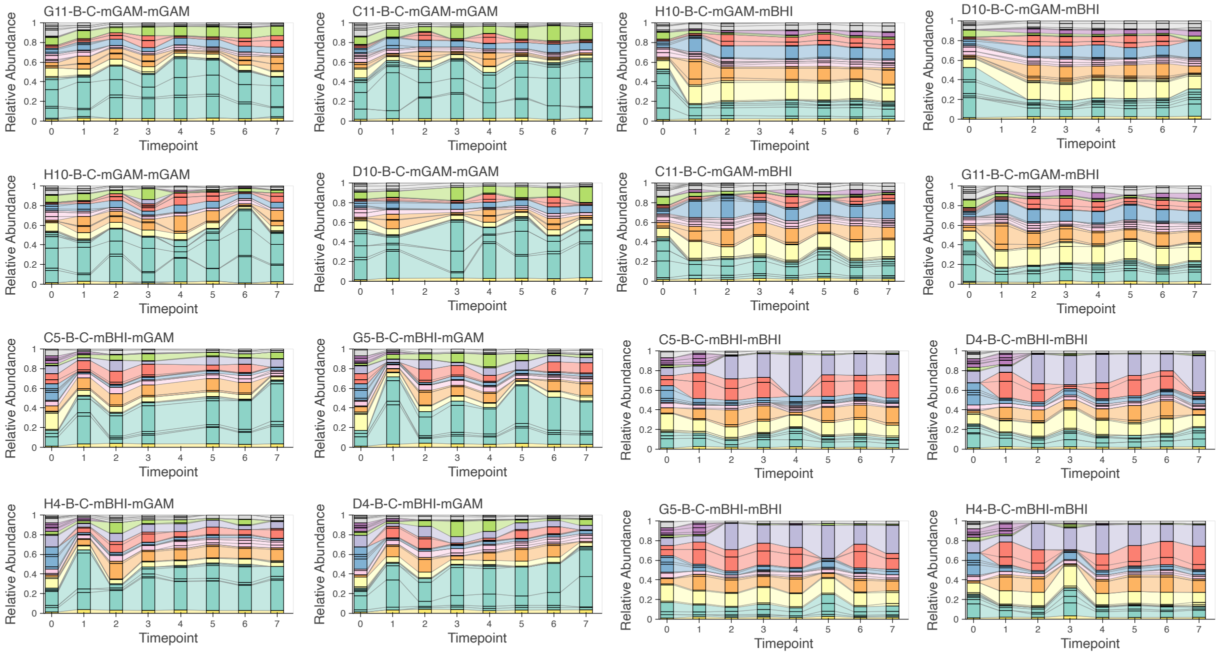


**Fig. S7: Analogous versions of Fig. 2A for all community collisions between hosts B and C.** Species are colored by bacterial family using the same color scheme as in Fig. 1.

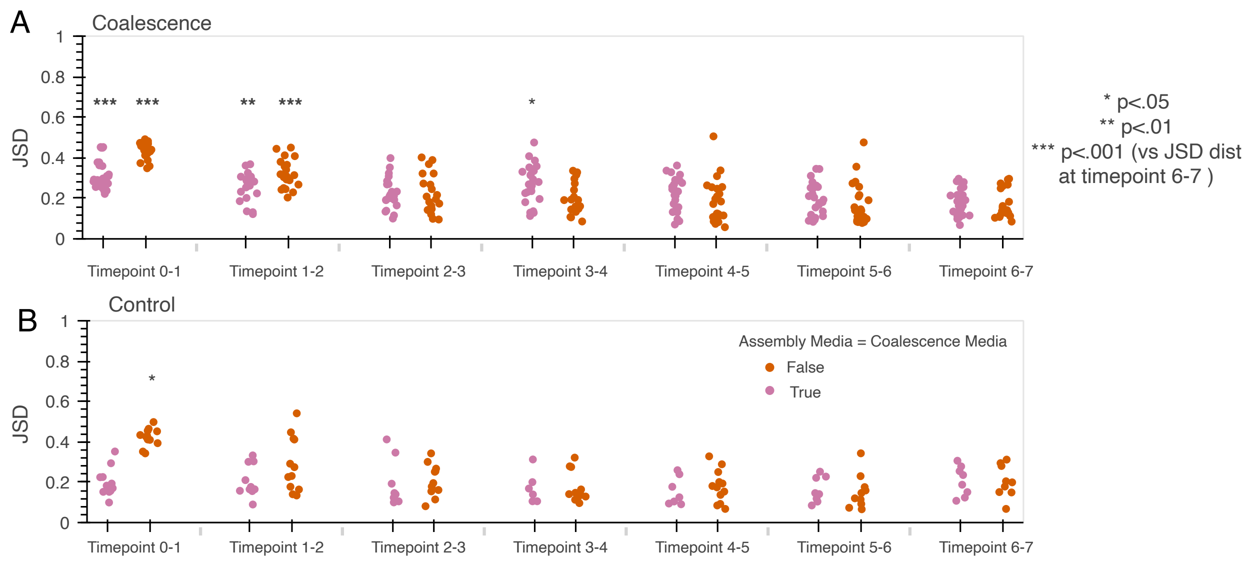


**Fig. S8: Quantifying the stability of community coalescence at the species-level using Jensen-Shannon distance.** Analogous versions of Fig. S2 for the coalescence (A) and control (B) arms of the community coalescence experiment. Points are colored by whether they underwent a media shift at timepoint 0. Timepoint 0 indicates the passage in which coalescence was performed, which is 7 dilution cycles after communities were derived from fecal samples. Taxonomic stability was evaluated by comparing the distribution of JSD values in each column to the JSD distribution between timepoints 6 and 7. Asterisks show significant permutation tests on the difference in median JSD values (***=p<0.001, **=p<0.01, *=p<0.05).

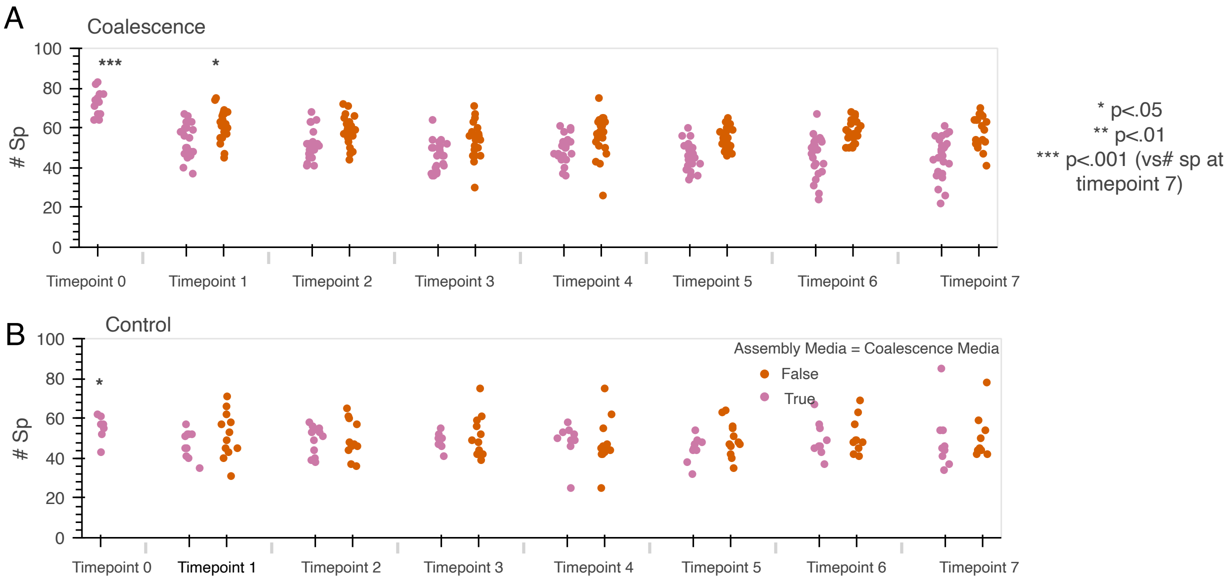


**Fig. S9: Quantifying the stability of community coalescence from the total species richness.** Expanded versions of Fig. 2B for the coalescence (A) and control (B) arms of the community coalescence experiment. As in Fig. S8, points are colored by whether they underwent a media shift at timepoint 0. Taxonomic stability was evaluated by comparing the richness distribution in each column to richness at passage 7. Asterisks show significant permutation tests on the difference in median richness values (***=p<0.001, **=p<0.01, *=p<0.05).

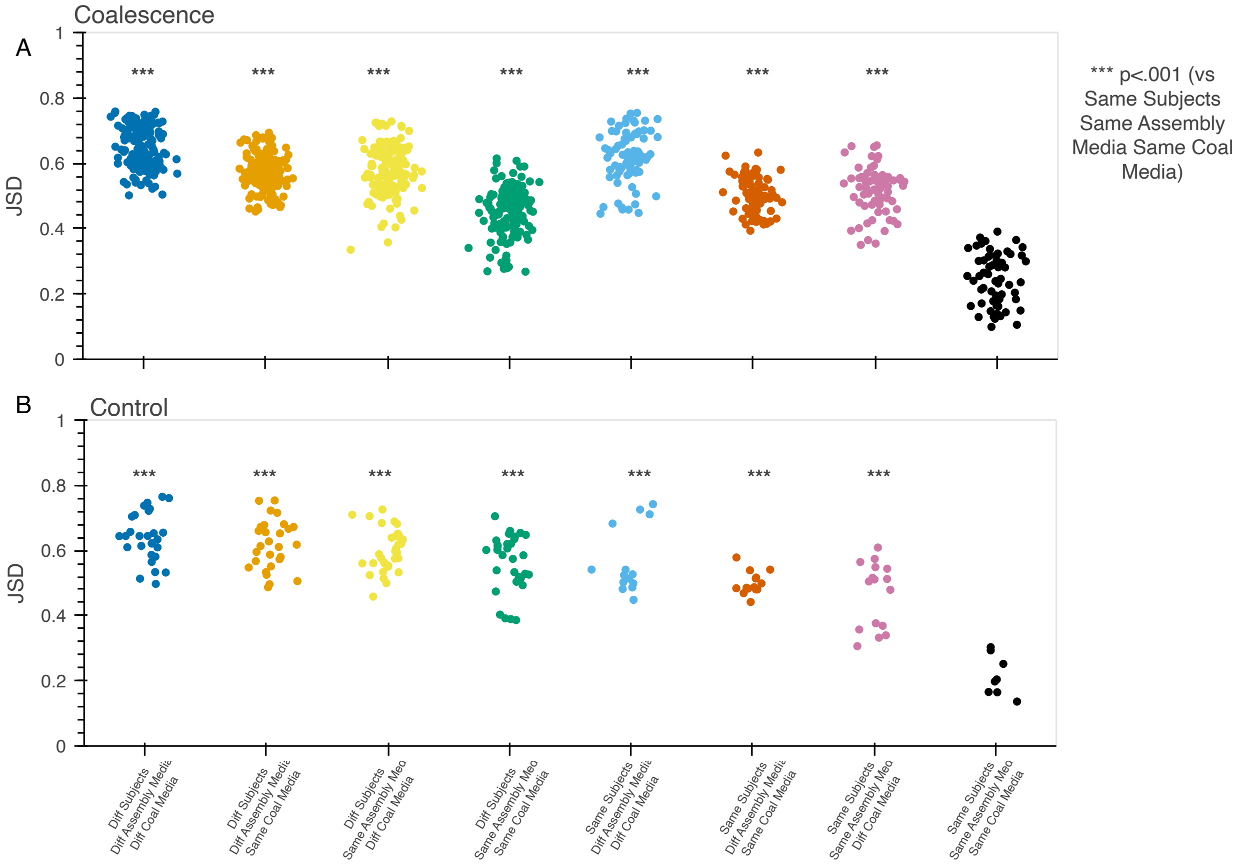


**Fig. S10: Community coalescence yields reproducible communities at the species level.** Between community JSD at timepoint 7 for different pairs of communities in the coalescence (A) and control (B) arms of the community coalescence experiment. For these comparisons, low-depth samples with fewer than 20 species were omitted. Reproducibility was evaluated by comparing the distribution of JSD values in each column to the biological replicates column on the right ('Same subjects Same Parent Media Same Coal Media'). Asterisks show significant permutation tests on the difference in median JSD values (***=$p<0.001$).

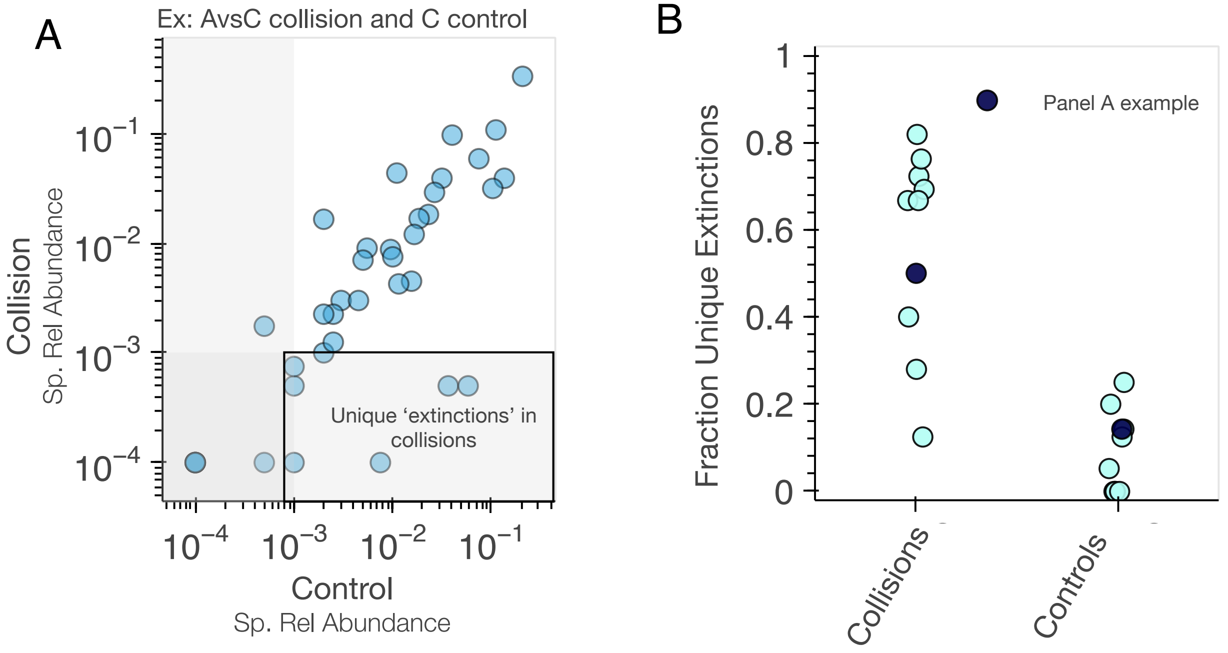


**Fig. S11: Community coalescence causes extensive species extinctions.** **(A)** Example data from the collision between hosts A and C (mGAM-mGAM). For each species that was present in host A at the time of the collision, points show the final mean relative abundance across replicate collisions as a function of their final mean relative abundance in the unmixed controls. Species were deemed to be "extinct" if their final relative abundance was $< 10^{-3}$. Points in the bottom right box represent species that went "extinct" in the collision but remained present in the unmixed control. **(B)** The fraction of extinction events that were unique to the collision (left) or control (right) communities for each of the $n = 12$ collisions. The example in panel A is highlighted in dark blue. The significant difference between these distributions (p<.001, permutation test, difference in medians) shows that many species that went extinct in the collision experiments did not go extinct in their respective controls.

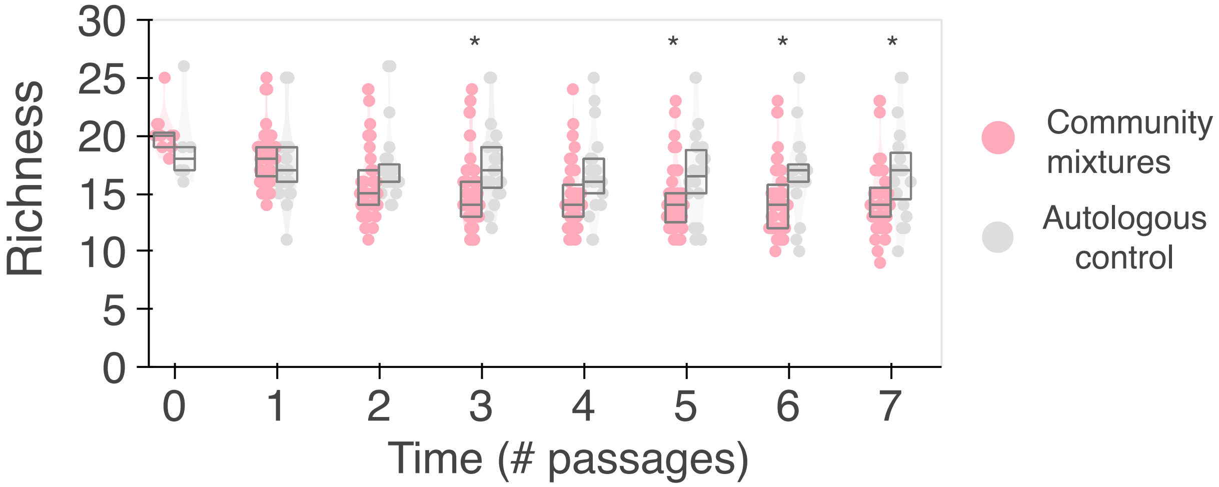


**Fig. S12: Community coalescence causes extensive species extinctions of higher abundance species.** Analogous version of Fig. 2B that only includes species with an initial relative abundance at passage 0 that was $\geq 1\%$. Many initially higher abundance species in collisions go extinct compared to those in controls, as seen by the fewer intiially higher abundance species present in collisions compared to control communities (permutation test of the difference in median between groups; *=P<0.05, **=P<0.01). Boxes show the median and inter-quartile range at each timepoint. These data show that the species that go extinct during community coalescence are not limited to rare taxa.

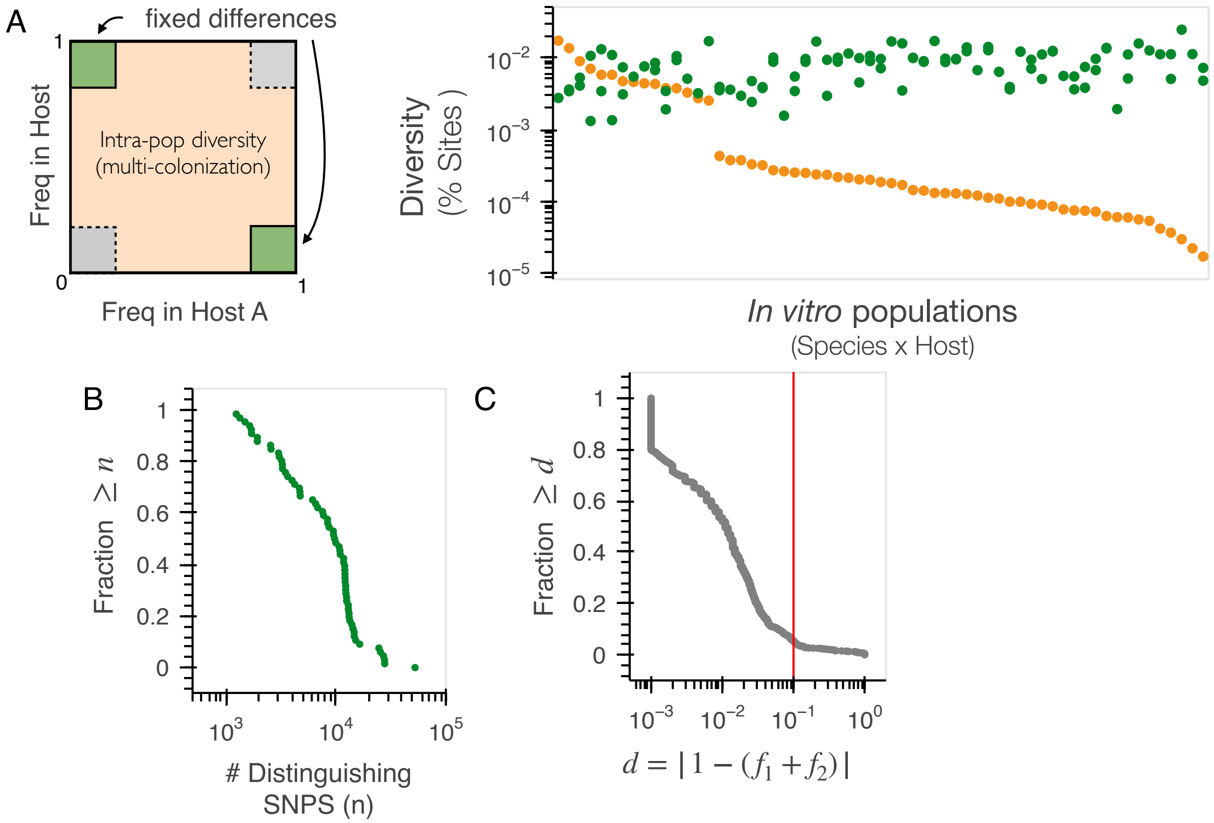


**Fig. S13: Inferring sets of marker SNVs to track conspecific strains in metagenomic data. (A)** Left: Schematic of how marker SNVs are inferred. SNVs in the dark green corners, which represent fixed or nearly-fixed differences between samples, are used to create two sets of marker SNVs (one from each corner; Methods). Right: analogous version of Fig. 1D showing within-sample diversity (orange) and between-sample divergence (green) for each species population in the samples used to inoculate a pairwise competition. **(B)** Distribution of the total number of identified marker SNVs for each pair of conspecific strains. **(C)** Distribution of the deviations of the two complementary marker SNV trajectories from one. The quantities $f_1$ and $f_2$ represent the two complementary strain frequencies estimated from the two independent sets of marker SNVs in panel A, which should theoretically sum to one. Large deviations from this expectation indicate potential inaccuracies in strain reconstruction. Red line indicates the threshold used to exclude samples from further consideration.

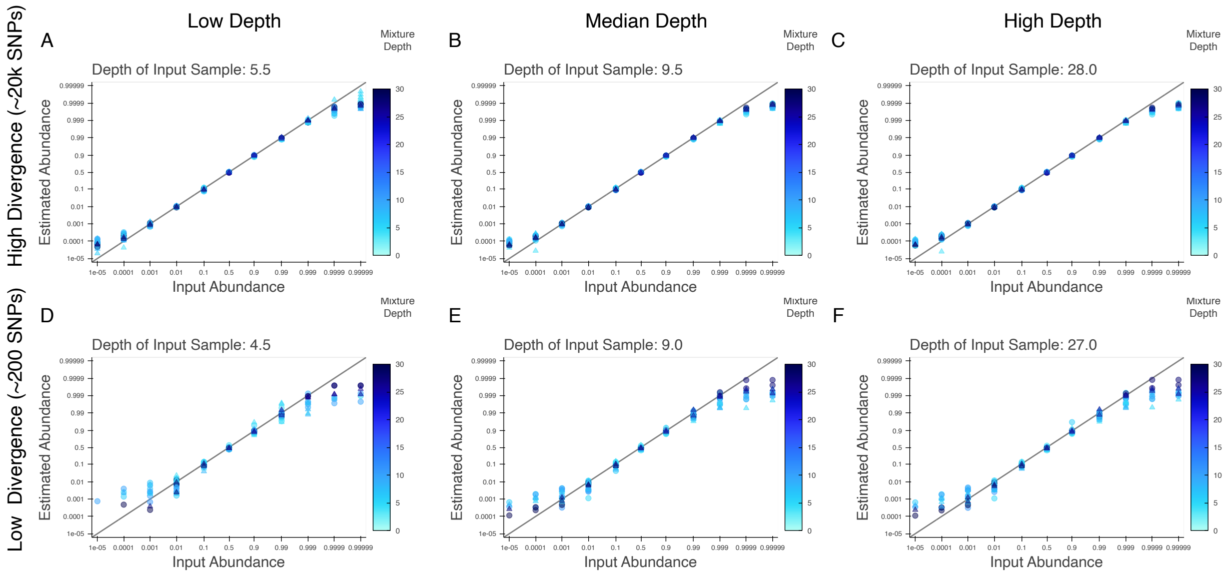


**Fig. S14: Validation of strain inference methods using simulated metagenomic data.** Each panel shows the estimated vs true relative strain frequencies in simulated metagenomes using a pair of typically related (A-C) and distantly related (D-F) *Bacteroides thetaiotaomicron* strains (Methods). Points are colored by the depth of the *B. theta* population in the mixture sample and differences in point shape indicate that different relative abundance profiles were used to generate samples for marker SNV inference (circle for A10-e003Coalescence-mBHI-p7, triangle for G8-e003Coalescence-mBHI-p7_S125). Plot titles indicate the depth of the parent samples that were used to identify marker SNVs.

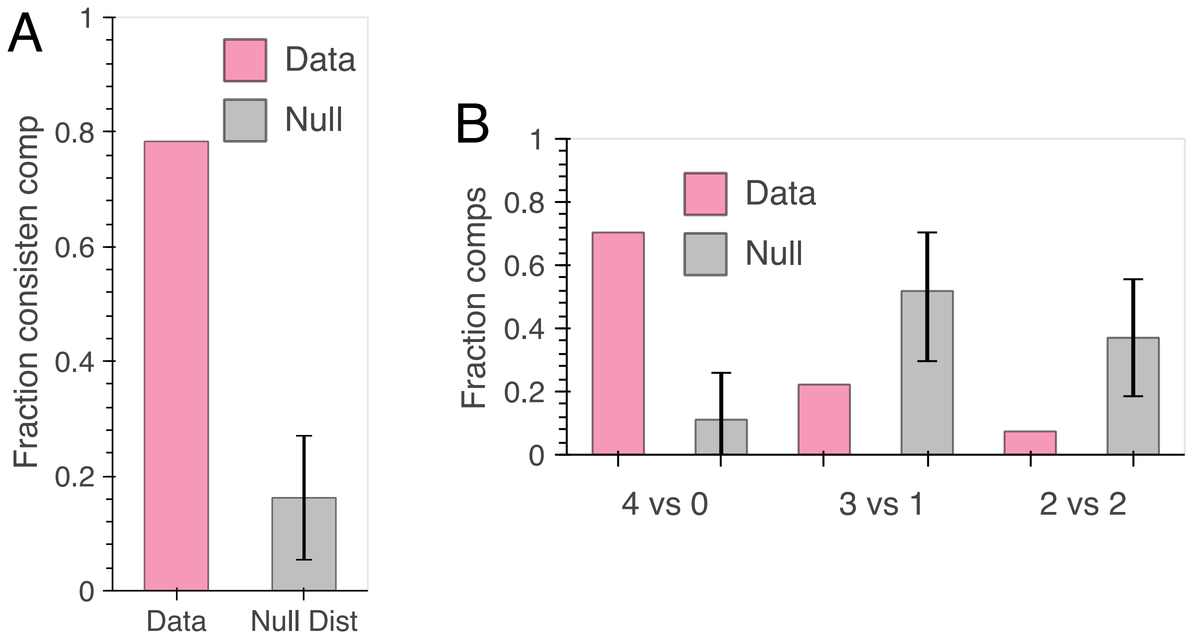


**Fig. S15: Fraction of strain competitions in which the direction of the temporal shift was consistent across replicates (A)** Fraction of strain competitions in which the direction of the temporal shift was consistent across all competitions with data from 3 or 4 replicates (n=37). Grey bars show the bootstrapped null distribution assuming that the direction of the shift is independent across replicates (error bars denote 95% confidence intervals). **(B)** Analogous version of panel A, but restricted to the subset of competitions with data from all four replicates (n=27). Outcomes are partitioned according to the number of independent replicates that shifted in one direction or the other (i.e. 4:0 indicates that all 4 replicates shifted in the same direction, while 2:2 indicates that 2 replicates shifted in one direction and 2 shifted in the other).

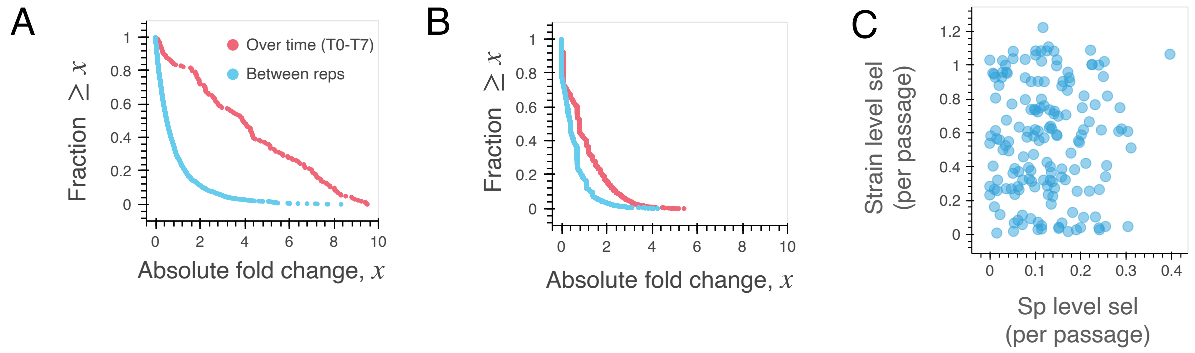


**Fig. S16: Comparing the strength of species- and strain-level selection (A)** Analogous version of Fig. 2F, replotted as a function of the absolute fold change between samples, $x = \log\left(f_i/(1-f_i)\right) \times ((1-f_j)/f_j))$. **(B)** Corresponding distribution of absolute fold changes at the species level, for all species with an initial relative abundance $\geq 10^{-3}$. **(C)** Scatter plot showing the effective species- and strain-level selection coefficients between passages 0-7 for all of the competitions in panel A. The correlation is between these two quantities is not statistically significant (Pearson's $\rho = 0.004$; $p > .05$)

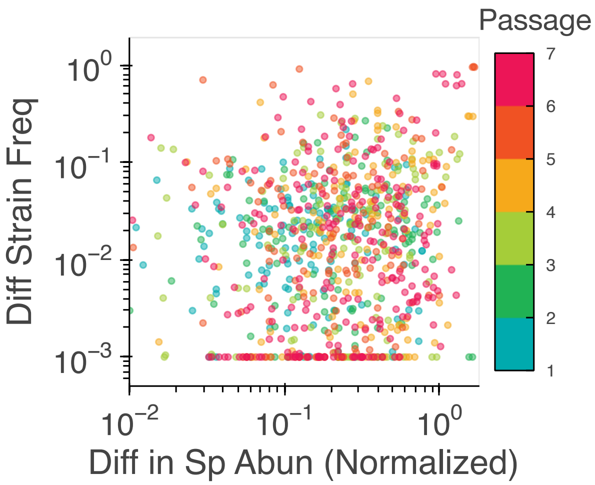


**Fig. S17: Relationship between the difference in conspecific strain frequencies the abundance of the focal species.** Each point shows a comparison between a pair of biological replicates in the community collision experiment, colored by the passage number. Differences in species abundance are scaled by the mean relative abundance of the focal species in the two replicate communities. There is a small but statistically significant correlation between these two quantities (Pearson's $\rho = 0.287$, $p < 0.001$).

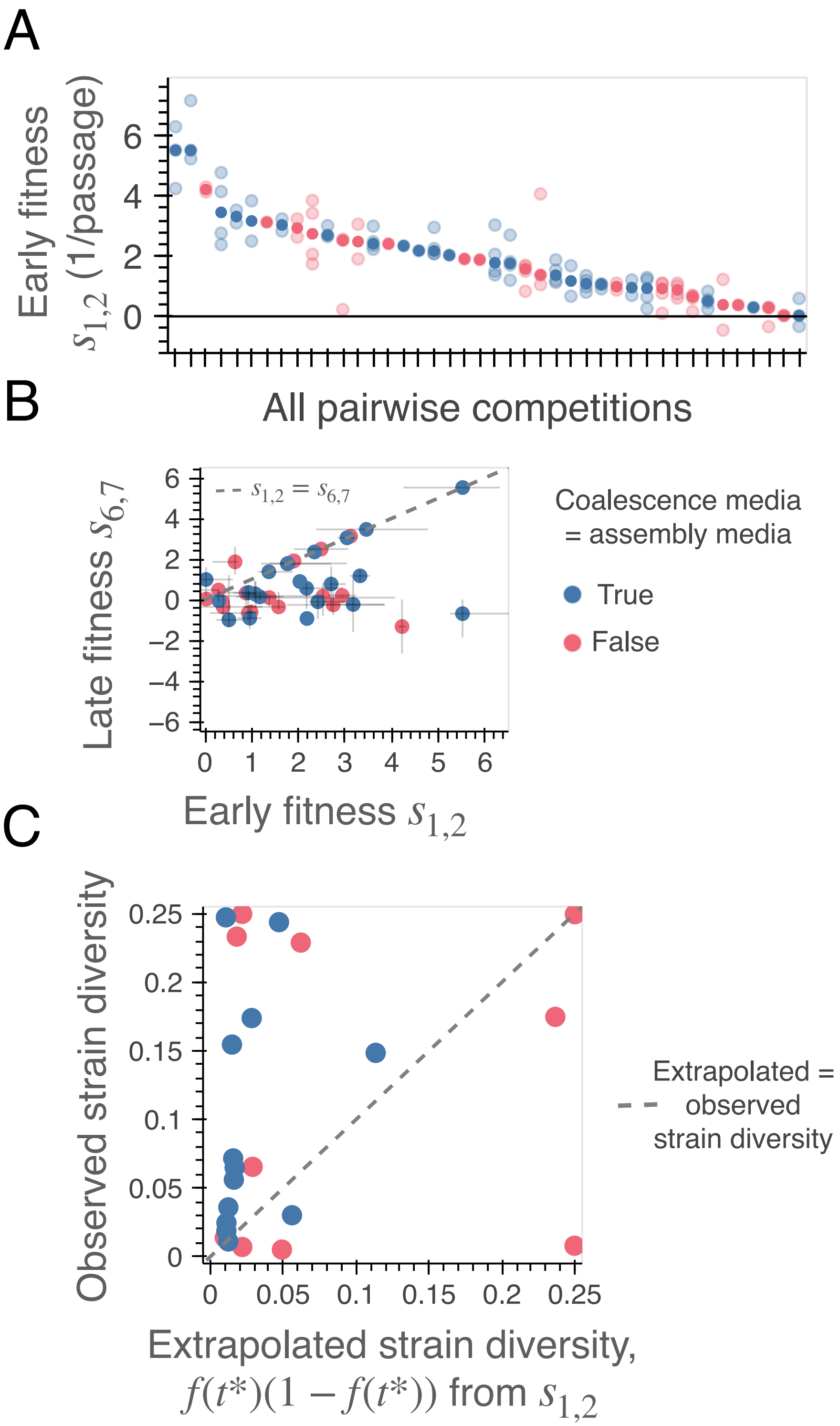


**Fig. S18: Relationship between strain-level selection and the effects of a media shift.** Analogous version of Fig. 3B-D in which each competition is colored by whether the collision was performed in the same or different growth media as the initial assembly. These data show that our observations of strong selection and intra-species coexistence are not restricted to collisions that have undergone a nutrient shift.

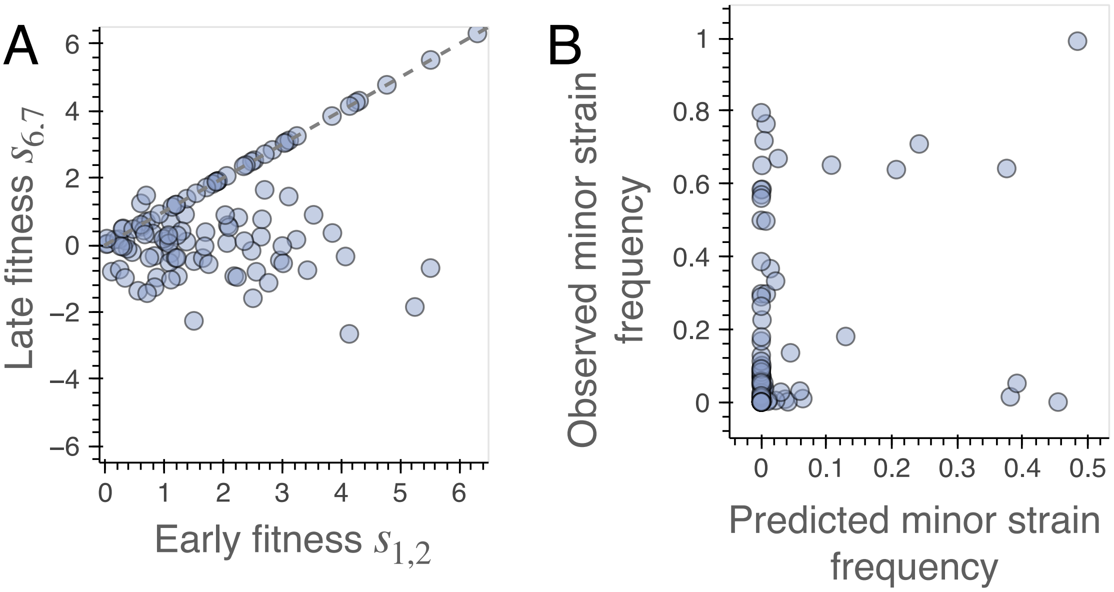


**Fig. S19: Analogous version of Fig. 3C showing all individual replicates. (A)** Estimated selection coefficients between passages 6 and 7 as a function of the selection coefficients between passages 1 and 2. As in Fig. 3C, we set $s_{6,7} = s_{1,2}$ for competitions in which the final frequency of the minor strain was below $5 \times 10^{-3}$. **(B)** Frequency of the predicted-to-be-minor strain at passage 7 versus the extrapolated frequency using the measured value of $s_{1,2}$.

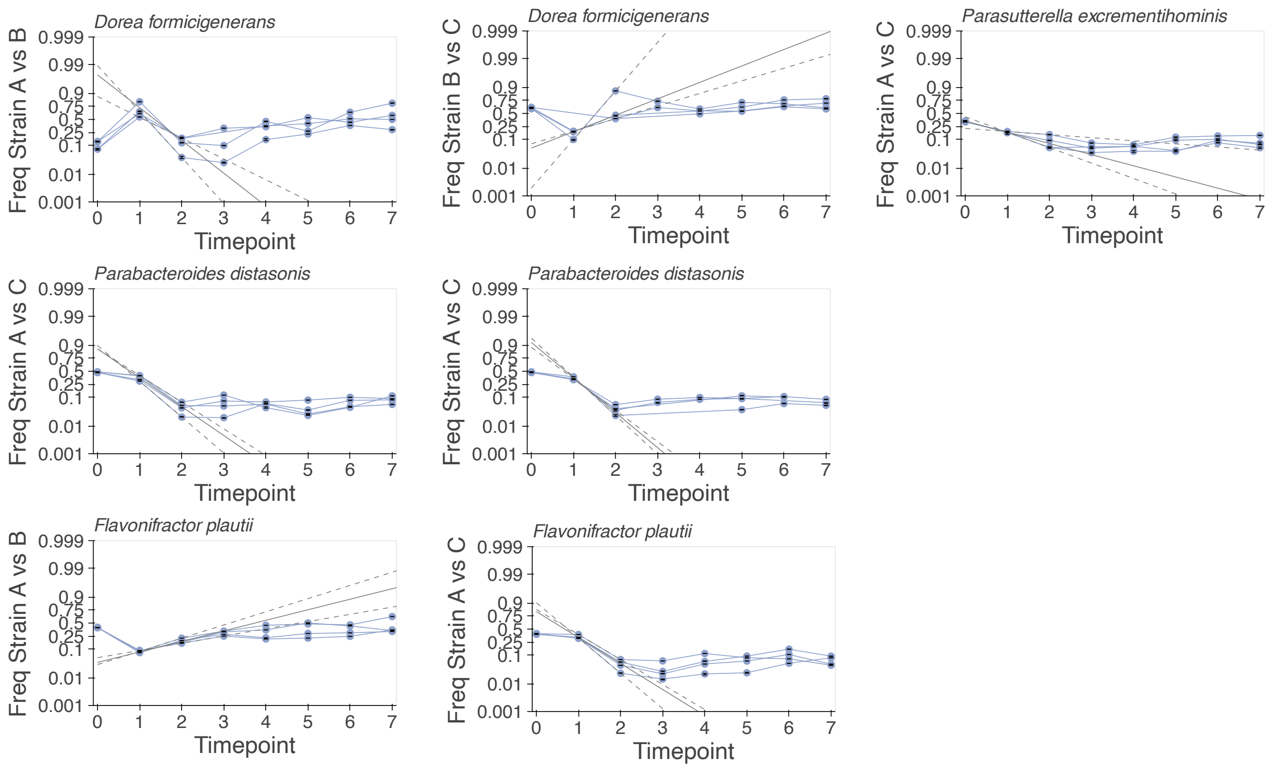


**Fig. S20: Additional examples of time-varying selection that were consistent across replicate collisions.** Analogous versions of Fig. 3A for the other species that exhibited consistent shifts in selection across replicates (q<0.05; t-test between $s_{1,2}$ and $s_{6,7}$ across biological replicates.

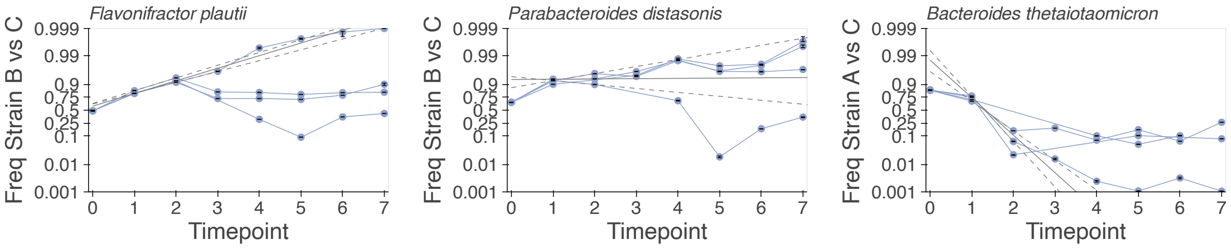


**Fig. S21: Examples of time-varying selection that were inconsistent across replicates.** Analogous versions of Fig. S20 for several examples where one or more of the replicates diverged from the others.

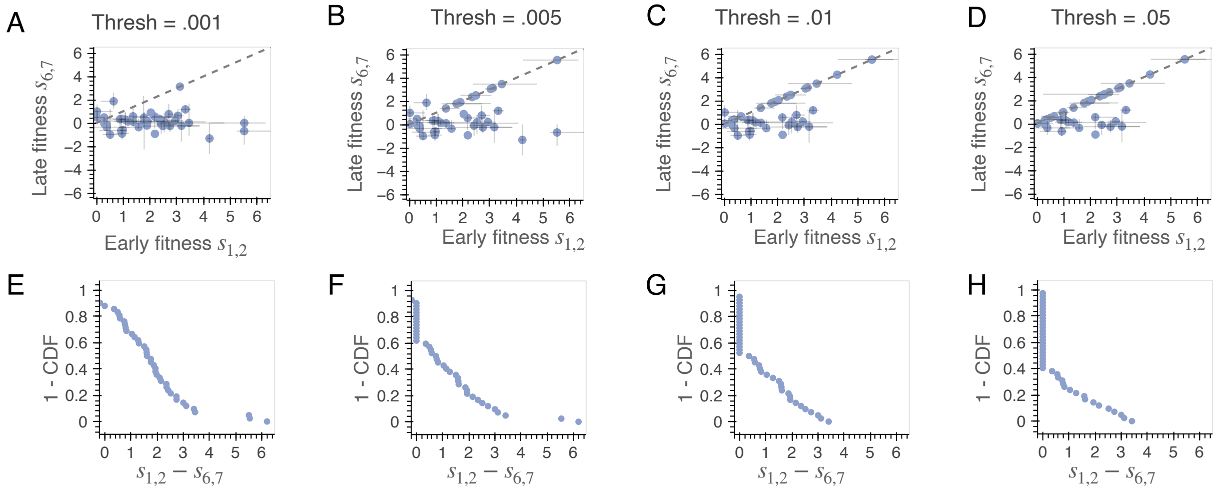


**Fig. S22: Attenuation of strain-level selection over time for different frequency cutoffs. (A-D)** Analogous versions of Fig. 3B with different frequency thresholds used for the resolution limit. As in Fig. 3B, if the median frequency of the minor strain across replicates fell below the indicated threshold then $s_{6,7}$ was set to be equal to $s_{1,2}$. (E-H) Distribution of the change in relative fitness ($s_{1,2} - s_{6,7}$) for the points in panels A-D.

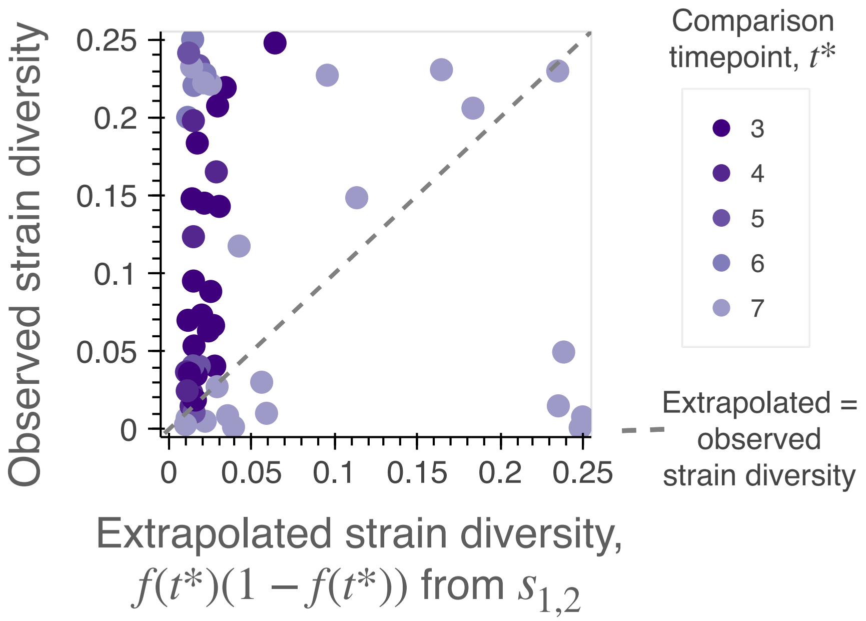


**Fig. S23: Analogous version of Fig. 3D showing all individual replicates**. The enrichment of points above the 1:1 line remains statistically significant (n=42/56; $p<0.01$; Methods).

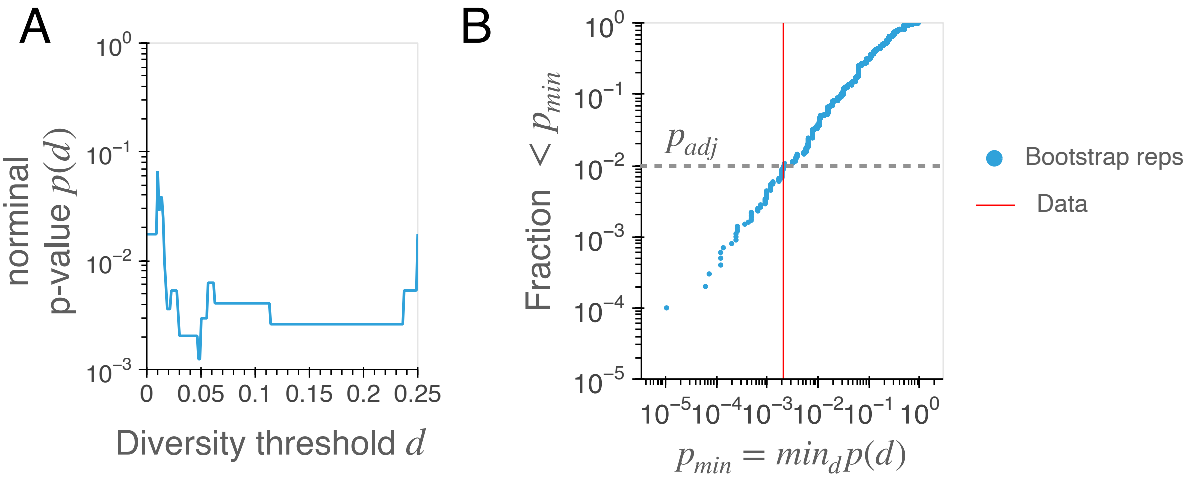


**Fig. S24: Statistical tests for assessing whether the shifts in $s(t)$ are biased in the direction of increasing intra-species diversity. (A)** For a given diversity threshold $d$, we can compute a nominal p-value $p(d)$ that quantifies whether the set of points in Fig. 3A with initial diversity $\leq d$ are more likely to fall above (versus below) the constant selection line (one sided binomial test, $p = 1/2$; Methods). These data show that the bias toward increased diversity is strongest among competitions with lower initial diversity ($d \approx 0.05$). **(B)** To account for multiple testing, we computed the distribution of the minimum value $p_{\min} \equiv \min_d p(d)$ across all thresholds for $n = 10,000$ bootstrap replicates of the original dataset (blue dots) and compared this distribution to the observed data (red). These data show that the adjusted p-value is $p_{\text{adj}} \approx 0.01$.

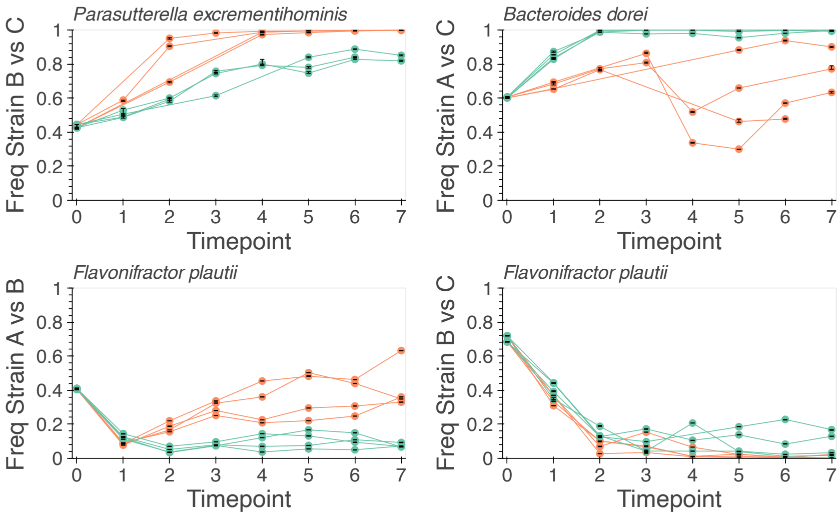


**Fig. S25: Additional examples of more gradual shifts in intra-species selection across abiotic environments.** Analogous versions of Fig. 4C for species with statistically significant selection trajectories in mBHI vs mGAM, using the same color scheme as Fig. 4.

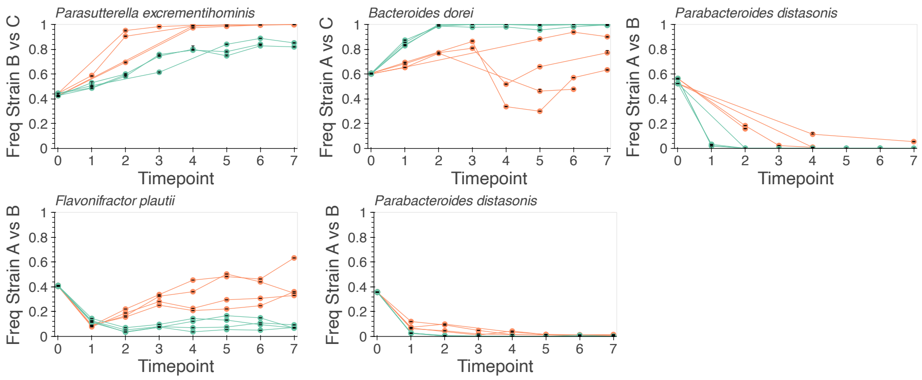


**Fig. S26: Additional examples of qualitative shifts in intra-species selection across abiotic environments.** Analogous versions of Fig. 4C for species with statistically significant selection trajectories in mBHI vs mGAM, using the same color scheme as Fig. 4.

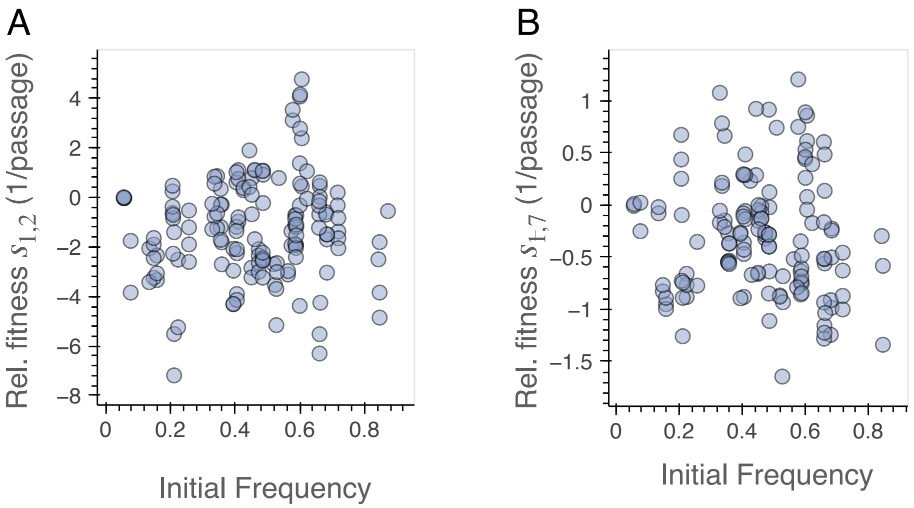


**Fig. S27: Relationship between initial relative abundance and competitive fitness of conspecific strains.** Points denote the estimated relative fitness of each strain pair as a function of their initial frequency, which reflects their relative abundance in the two parent communities immediately prior to mixing. Left panel (A) shows the estimated relative fitness in the early phase of the competition (Fig. 3 B), while the right panel (B) shows the net relative fitness between passages 1 and 7. Neither variable exhibits a statistically significant correlation with the initial strain abundance (Pearson's $\rho = 0.095, p = 0.242$ for panel A and $\rho = -0.0843, p = 0.322$ for panel B).

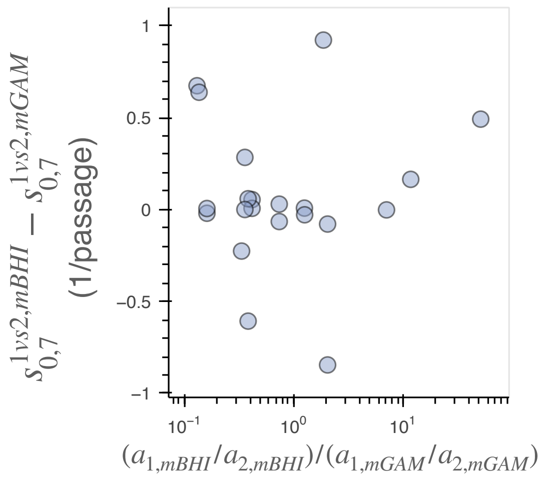


**Fig. S28: Relationship between the shift in relative abundance and shift in competitive fitness across media.** Analogous version of Fig. S27 comparing the shift in relative fitness between the two growth media (Fig. 4B) with the shift in relative abundance ($a$) of the two conspecific strains in their corresponding parent communities. The correlation between these two quantities is not statistically significant (Pearson's $\rho = 0.240, p = 0.300$).

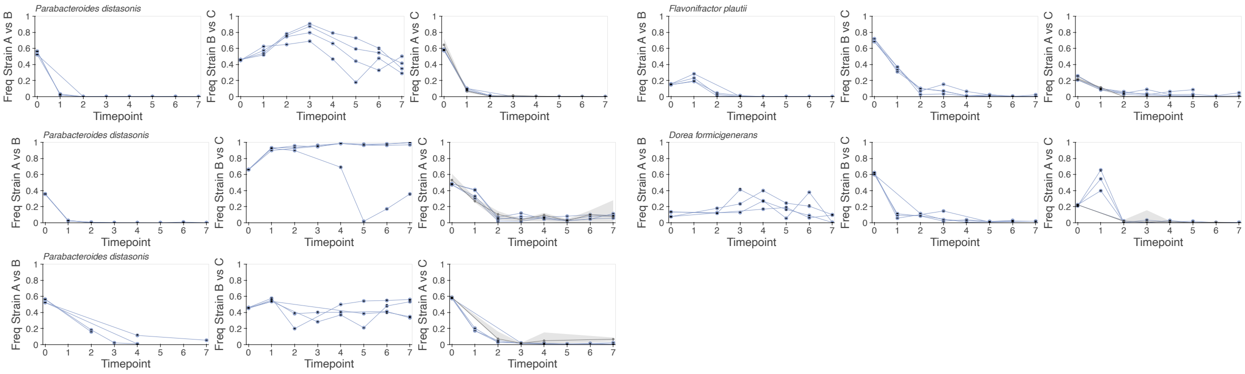


**Fig. S29: Additional examples of transitive competition trios.** Analogous versions of Fig. 5A for other example species that were consistent with the transitive fitness model (Methods).

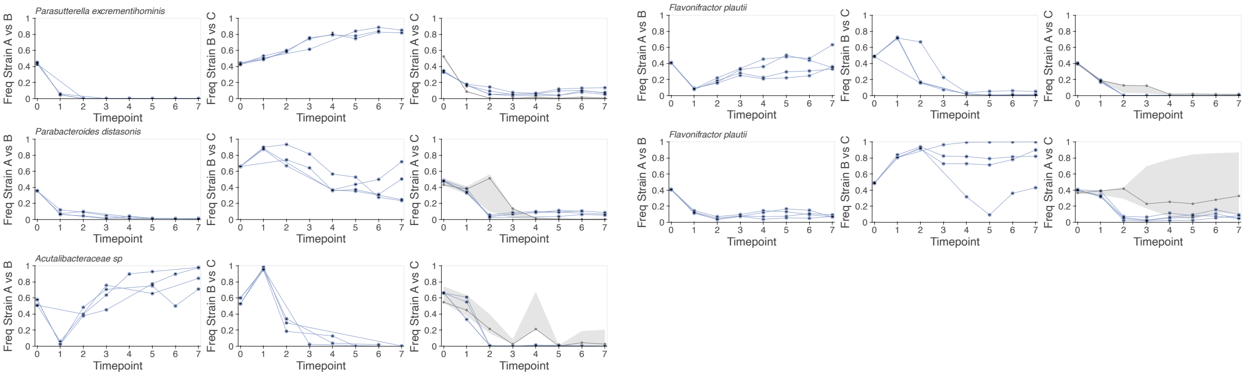


**Fig. S30: Additional examples of non-transitive competition trios.** Analogous versions of Fig. 5A for other example species that exhibited deviations from the transitive fitness model (Methods).

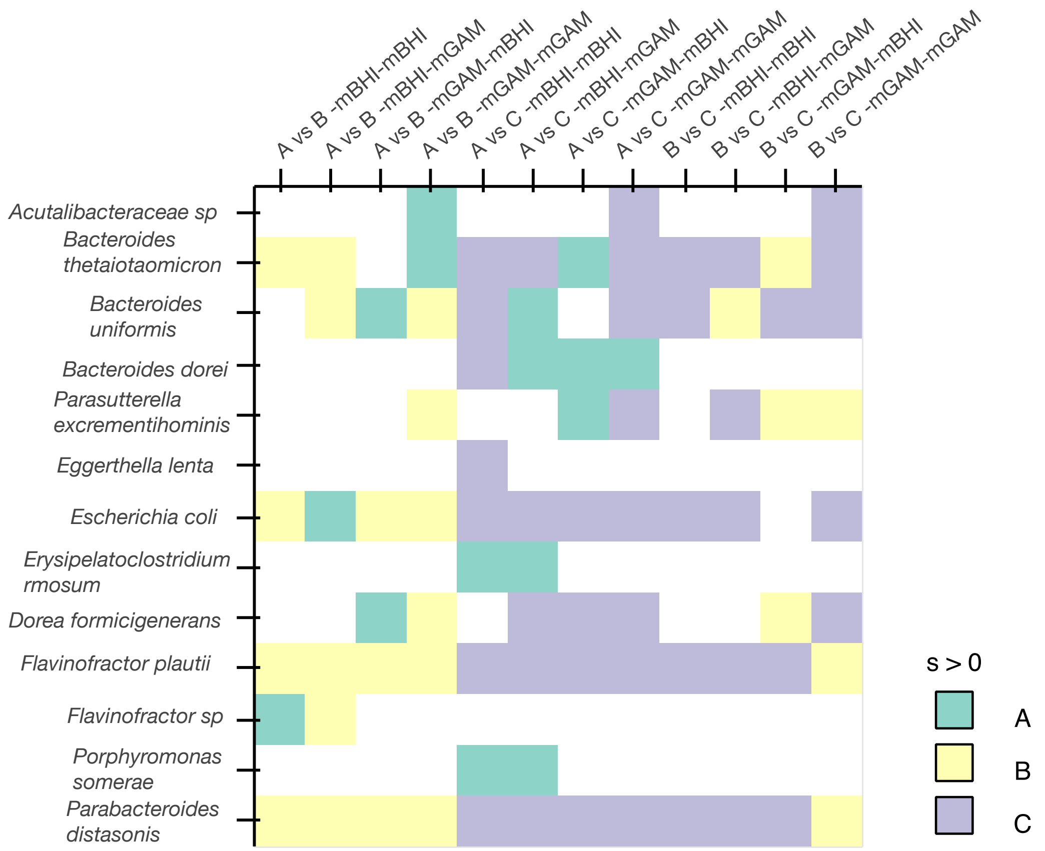


**Fig. S31: Analogous version of Fig. 5C showing the outcomes for individual species.** Columns illustrate different pairwise coalescence experiments, while entries are colored by the host in which the median net selection coefficient between passages 0 and 7 was positive. White boxes indicate cases where the species was absent (or at low coverage) for that competition. These data show that most collisions exhibited a mixture of winning strains from both hosts, and that most species exhibited a range of outcomes across collisions.

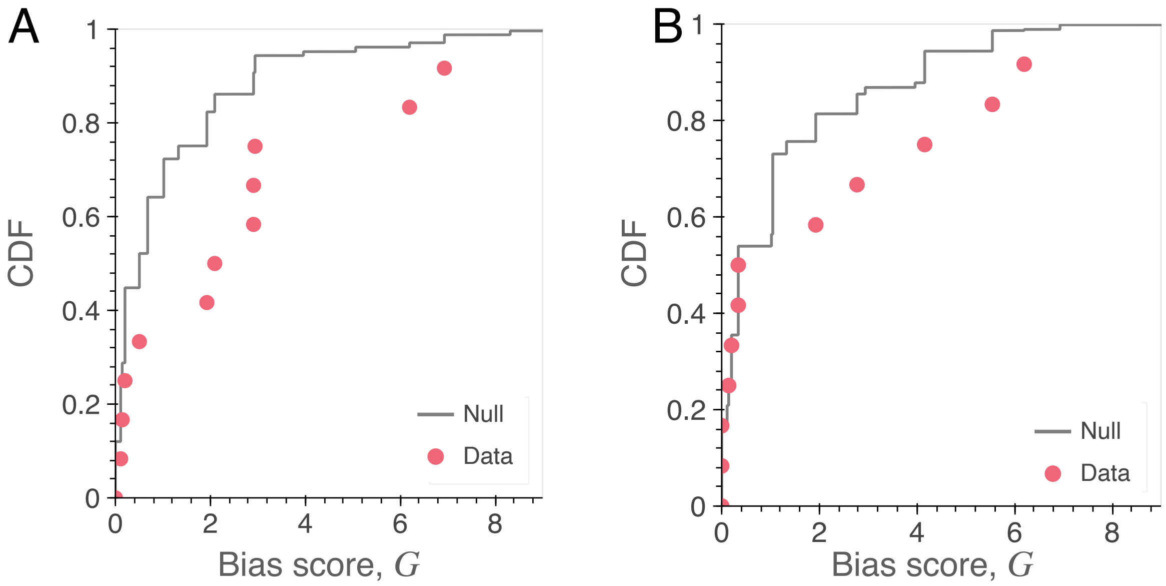


**Fig. S32: Statistical tests for strain-level community cohesion. (A)** Distribution of the bias score $G$ (Eq. 9) for all $n = 12$ competitions (points) compared a null model where the winning strains were randomly from each host with equal probability (line). The observed distribution is different from the null distribution (median observed host bias = 2.01, null host bias = 0.51, p=0.002; Methods). **(B)** Analogous version of panel A, but restricted to competitions where species were dominated by one strain. In this case, the observed bias distribution is not significantly different from the null (p>0.05)

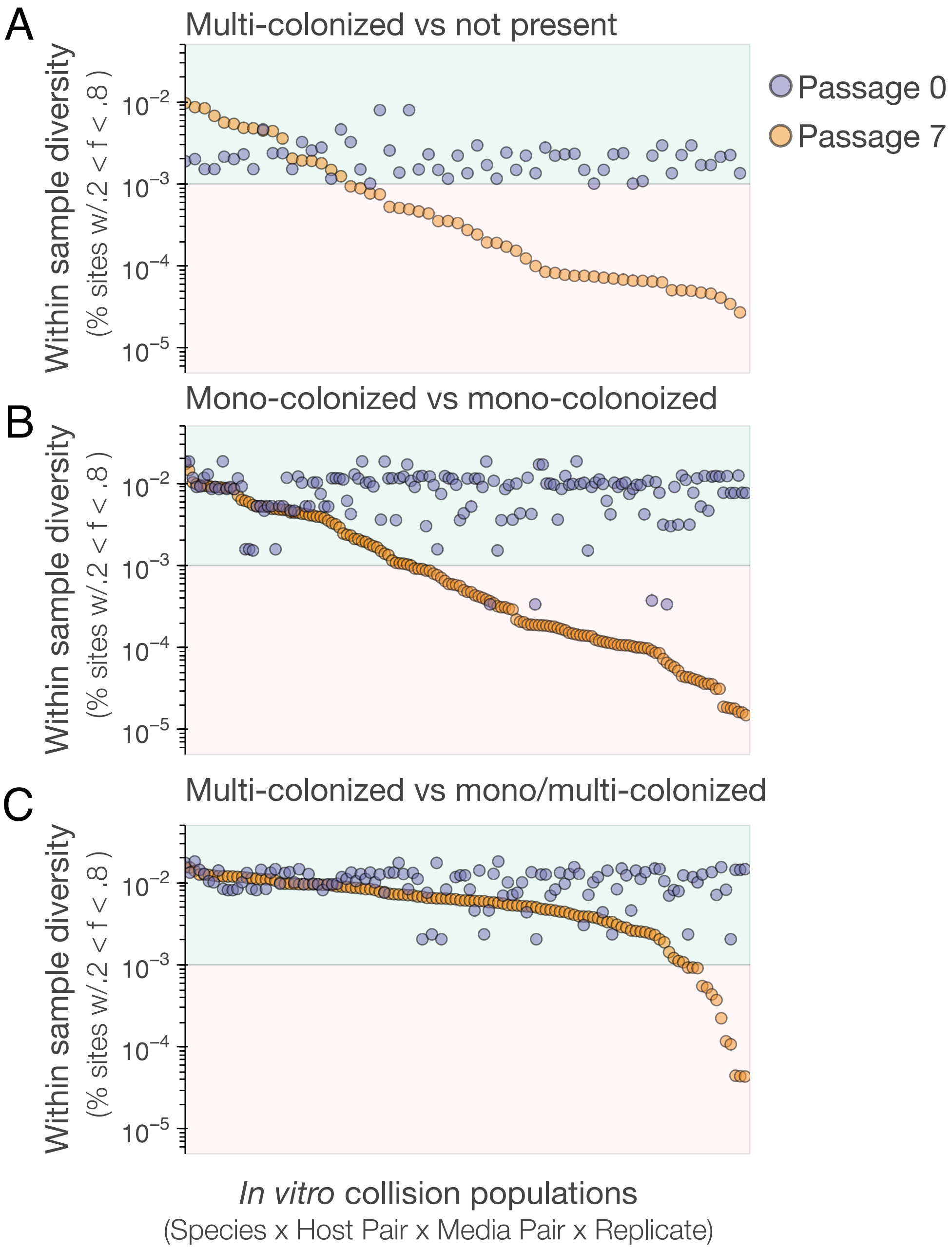


**Fig. S33: Signatures of multi-colonization during community coalescence. (A-C)** Analogous versions of Fig. 1D constructed for the community collision experiments. Points are colored by whether they are from timepoint 0 or timepoint 7 post coalescence. Panel A shows all populations that were initially multi-colonized in one parent and not present in the other host, panel B shows populations that were initially mono-colonized in the parent communities from each host, and panel C shows populations that were present in both parent communities and were initially multi-colonized in at least one parent community. These data show that intra-species coexistence is not limited to strains with a prior history of co-occurence.

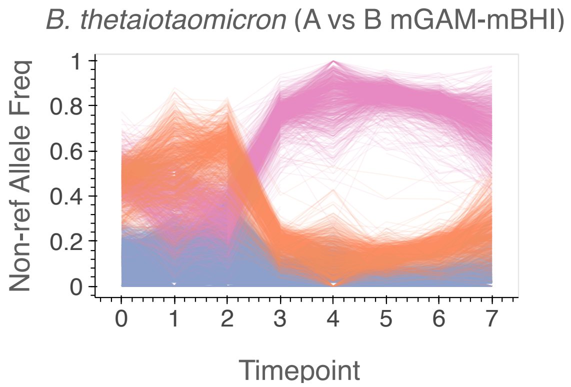


**Fig. S34: Example of SNV trajectories used for tracking major strains from multi-colonized populations.** In this example, the population from host B prior to collision had elevated levels of genetic diversity (1.3% of sites with intermediate frequency SNVs; Fig. 1D)) while the population from host A was mono-dominated (0.008% of sites with intermediate frequency SNVs). Marker SNVs for the host A strain are colored pink, marker SNVs for the major strain from host B are colored orange, and all other intermediate-frequency SNVs are colored lavender. In this competition, the strain from host A rises to high frequency to dominate the mixed population. The major strain from host B travels to lower frequency, and SNVs from the minor strain(s) of host B (lavender) fall to lower frequencies as well.

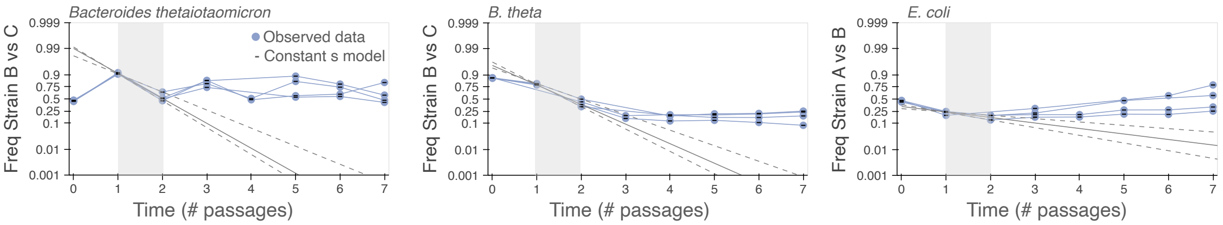


**Fig. S35: Examples of shifting selection pressures in multi-colonized species.** Analogous versions of Fig. 3A for several competitions which had at least one multi-colonized parent population.

**Table S1:** A list of all samples, accessions, and total sequencing depth for the metagenomic data generated in this study.

**Table S2:** Demographic information for all study participants who provided the initial stool samples.

**Table S3:** Sample metadata for the community assembly portion of the study.

**Table S4:** Sample metadata for the community coalescence portion of the study.

**Table S5:** List of all strain pairs for which competition could be assessed during community coalescence, along with their corresponding colonization status.
